# Region-specific, maladaptive, gray matter myelination is associated with differential susceptibility to stress-induced behavior in rats and humans

**DOI:** 10.1101/2021.02.15.431176

**Authors:** Kimberly L. P. Long, Linda L. Chao, Yurika Kazama, Anjile An, Kelsey Y. Hu, Lior Peretz, Dyana C. Y. Muller, Vivian D. Roan, Rhea Misra, Claire E. Toth, Jocelyn M. Breton, William Casazza, Sara Mostafavi, Bertrand R. Huber, Steven H. Woodward, Thomas C. Neylan, Daniela Kaufer

**Author notes:** Current address: Department of Psychiatry and Behavioral Sciences, University of California, San Francisco, San Francisco, CA 94143, USA. Current address: Department of Psychiatry, Columbia University, New York, NY, 10027, USA.

## Abstract

Individual reactions to traumatic stress vary dramatically, yet the biological basis of this variation remains poorly understood. Recent studies demonstrate the surprising plasticity of oligodendrocytes and myelin with stress and experience, providing a potential mechanism by which trauma induces aberrant structural and functional changes in the adult brain. In this study, we utilized a translational approach to test the hypothesis that gray matter myelin contributes to traumatic-stress-induced behavioral variation in both rats and humans. We exposed adult, male rats to a single, severe stressor and used a multimodal approach to characterize avoidance, startle, and fear-learning behavior, as well as oligodendrocyte and myelin content in multiple brain areas. We found that oligodendrocyte cell density and myelin content were correlated with behavioral outcomes in a region-specific manner. Specifically, stress-induced avoidance positively correlated with hippocampal dentate gyrus oligodendrocytes and myelin. Viral overexpression of the oligodendrogenic factor Olig1 in the dentate gyrus was sufficient to induce an anxiety-like behavioral phenotype. In contrast, contextual fear learning positively correlated with myelin in the amygdala and spatial processing regions of the hippocampus. In a group of trauma-exposed US veterans, T1-/T2-weighted magnetic resonance imaging estimates of hippocampal and amygdala myelin associated with symptom profiles in a region-specific manner that mirrored the findings in rats. These results demonstrate a species-independent relationship between region-specific, gray matter oligodendrocytes and myelin and differential behavioral phenotypes following traumatic stress exposure. This study suggests a novel mechanism for brain plasticity that underlies individual variance in sensitivity to traumatic stress.

## Introduction

Exposure to a traumatic stressor induces a spectrum of outcomes in both humans and animals, ranging from no observable changes in behavior to long-lasting changes across various behavioral domains (1–5). For example, persistent, trauma-induced changes to avoidance-, fear-, and arousal-related behaviors define the psychiatric criteria for anxiety or post-traumatic stress disorder (PTSD) in humans (6, 7); however, not all who experience trauma develop subsequent mood and anxiety disorders (2–5). The underlying physiological mechanisms of this selective susceptibility and resilience to trauma are poorly understood. Furthermore, biomarkers of susceptibility along with novel therapeutic targets for anxiety and PTSD are lacking.

Recent studies have uncovered a surprising role for oligodendrocytes and the myelin they produce in stress, behavior, and experience-dependent plasticity (8–13). Namely, neuronal activity and hormonal signaling can induce the proliferation of oligodendrocyte precursor cells and the remodeling of existing myelin in regions such as the corpus callosum, somatosensory cortex, and premotor cortex (14–19). Enhancing or inhibiting this plasticity alters behaviors such as motor function, spatial memory consolidation, and motor learning (14,20,21), suggesting that adult myelin plasticity in both white and gray matter regions is necessary for proper synaptogenesis, circuit function, and learning. In addition, oligodendrocytes express glucocorticoid receptors throughout their lineage and are sensitive to various environmental stressors in a region-specific manner (22–28). For example, chronic stress decreases myelin density in the prefrontal cortex (PFC) (13,29,30), while early weaning in mice increases myelin production in the amygdala (31). In addition, chronic stress or exposure to stress-level glucocorticoids in adult, male rats increases oligodendrogenesis from neural stem cells in the dentate gyrus of the hippocampus (32).

Oligodendrocyte and myelin plasticity may ultimately contribute to fear learning and affective behavior in both rodents and humans (33–44). For example, conditional ablation of oligodendrocyte precursors in the PFC increases anxiety-like behavior and decreases social behavior in mice (45). Inhibiting the formation of new myelin in mice impairs remote fear memory recall (26). In humans, early life stressors such as institutionalization and neglect alter white matter microstructure in the PFC, and these alterations are correlated with PFC-related cognitive deficits (37). Patients with major depressive disorder display changes to white matter in the frontal-limbic system that correlate with behavioral symptoms such as rumination (39,40,44). Recent evidence has also raised the intriguing possibility that myelination in gray matter regions such as the amygdala and hippocampus plays a role in psychiatric disorders (42, 46). For example, patients with major depressive disorder display decreased oligodendrocyte cell density in the amygdala (42). Furthermore, we have shown that patients diagnosed with PTSD have significantly higher myelin content in the hippocampus as compared to trauma-exposed subjects without PTSD, and hippocampal myelin levels positively correlate with PTSD symptom severity (46). This suggests that oligodendrocytes and myelin in gray matter regions may correspond to individual differences in trauma-induced psychiatric pathology.

The sensitivity of oligodendrocytes to neuronal activity and stress, coupled with the alterations to oligodendrocytes and myelin seen in various psychiatric disorders, suggests that oligodendrocytes and myelination may serve as a mechanism by which stress produces adverse psychiatric outcomes (10, 47). The current study tests this concept in a translational framework. First, we utilized a multivariate approach to characterize the variability of individual and domain-specific components of behavior in a rat model of exposure to a single, acute trauma. Next, we performed microstructural analyses of oligodendrogenesis in the hippocampus, amygdala, and corpus callosum in relation to quantitative measures of avoidance, startle, and contextual fear memory. We then conducted Olig1-driven gain of function in the hippocampus to test whether altering oligodendrocyte content within the hippocampus is sufficient to induce a behavioral phenotype in the absence of stress exposure. In humans, we extended our analysis of myelin in the brains of US veterans (46) by including the amygdala and analyzing domain-specific PTSD symptom data.

## Materials and Methods

### Study design - Rodents

The aim of this study was to determine the relationship between oligodendrocytes, myelin, and the behavioral outcomes of trauma. Rats were exposed to an acute, severe stressor, and outcomes were assessed using behavioral (avoidance, fear, and startle assays) and molecular (immunofluorescent staining) measures. An experimental gain-of-function (Olig1 upregulation) was used to investigate whether oligodendrogenesis causally contributes to behavioral outcomes. Complementary measures of myelin in human subjects (via T1-/T2-weighted magnetic resonance imaging) were compared to sub-components of the Clinician Administered PTSD Scale (CAPS). The experimental designs and methodology of outcome measures were chosen prior to initiating each experiment. For all animal experiments, subjects were randomly assigned to experimental groups. Investigators were blind to stress and virus conditions during behavioral assays and data quantification. The criteria for animal exclusion were surgical complications and unsuccessful viral targeting. Study design and sample sizes used for each experiment are provided in figure legends.

### Animals

All animal care and procedures were approved by the University of California, Berkeley, Institutional Animal Care and Use Committee. A total of 70 adult (postnatal day 65) male Sprague Dawley rats were purchased from Charles River and pair-housed on a 12:12 light-dark cycle (lights on at 0700 hours) in our facility at UC Berkeley. Rats had *ad libitum* access to food and water and were given at least one week to acclimate to the facility before testing began. Rats underwent gentle handling (being picked up and held) for 5 days immediately prior to stress. Animals were weighed before and periodically following stress. For the first study, 40 male animals were used (20 control and 20 stress). These numbers were based on previous work with acute stress and behavior profiling in rats (48). For the fear conditioning study, 12 males were used. This number was based on a power calculation using the correlations detected in the dentate gyrus. Alpha was set to 0.05 and power to 80%, and the resulting necessary number ranged from 9 to 19 animals. Finally, for the viral work, 18 male rats were used (10 control virus and 8 Olig1 virus). This number was based on a power calculation using our GSTπ and BrdU data, which yielded a necessary number ranging from 2 to 6 animals per group.

### Stress

To establish a rodent model with which to study acute, severe trauma exposure, we coupled a single exposure to predator scent (fox urine) with 3 hours of immobilization in adult, male Sprague-Dawley rats. Rats were randomly assigned to undergo stress or remain in the home cage with minimal disturbance (BrdU injection and cage change, detailed below). Rats were run in pairs in cohorts of 10 animals each, with numbers of control and stress animals counterbalanced between cohorts. Rats exposed to stress underwent acute immobilization with exposure to predator scent. Cagemates were placed side-by-side in an empty cage inside a fume hood from 0900-1200 hours while being restrained in Decapicone bags (Braintree Scientific, Braintree, MA). A cotton ball infused with 1 mL of fox urine (Trap Shack Co. Red Fox Urine, amazon.com) was placed in the center of the cage. Tail vein blood was collected from stress-exposed rats at 0 minutes, 30 minutes, and 3 hours into acute immobilization stress. After cessation of stress, animals were injected with BrdU (detailed below) and were released and returned to a clean cage to allow for self-grooming. After 1 hour, rats were returned to a clean home cage. In order to minimize the transfer of stress and fox odors to the colony, stress-exposed animals were kept in a separate housing room for three days prior to being returned to the normal housing room. Rats in the control group received BrdU injections and a cage change at the same time of day.

### Serum corticosterone

At 0 minutes, 30 minutes, and 3 hours into acute immobilization stress, tail vein blood was collected from each rat for corticosterone sampling. Blood samples were centrifuged at 9,391 g for 20 minutes at 4°C, and serum was extracted and stored at -80°C. Samples were assayed using a Corticosterone EIA kit (Arbor Assays, Ann Arbor, MI). Any sample running below the detection limit of the assay was assigned the value of the detection limit (15.625 ng/mL).

### BrdU injections

To quantify oligodendrogenesis and/or oligodendrocyte survival following acute stress exposure, rats were injected intraperitoneally with 100 mg/kg 5-Bromo-2′-deoxyuridine (BrdU, Sigma-Aldrich, St. Louis, MI) dissolved in warm 0.9% saline immediately after release from Decapicone bags. They were then injected once each day for the following two days within one hour of the circadian time of the first injection. Control animals were injected at similar times of the day, taking care to avoid transfer of fox and stress odors.

### Behavioral battery

Animals underwent a behavioral battery 7 days after stress exposure to characterize the extent of persistent behavioral changes. All behaviors were conducted between 0800-1400 hours. Prior to all tests, animals were brought to the testing space and allowed at least 30 min to acclimate. One day prior to stress, all animals went through a 5 min baseline open field test (OFT) under dim lighting (15 lux). Seven days after acute immobilization stress, all rats were individually profiled for anxiety-like behaviors using 6 different behavioral tests: OFT in a brightly lit environment (OFT Light), OFT in a dimly lit environment (OFT Dim), light/dark box (LD), elevated plus maze (EPM) in a bright environment, EPM in a dim environment, and acoustic startle response (ASR). Low light versions of the OFT and EPM were utilized as low anxiogenic versions of their full light counterparts, allowing for further characterization of exploratory and anxiety-like behavior. The battery spanned two days with the following sequence: Day 1 - - OFT Light, EPM Light, ASR; Day 2 – OFT Dim, LD, EPM Dim. After placing an animal into an arena, the experimenter exited the room. Animals were given 10 minutes of rest in the home cage in between each test. Of note, our behavior facility underwent construction over the course of this study, and our apparatuses were moved to a new suite after experiment 1 (corresponding to figures 1 and 2). Behavior tests for virus experiments (corresponding to figure 3) were conducted in new rooms under identical conditions, except for the EPM Dim, in which dim red light was substituted for dim white light.

**Fig. 1:**
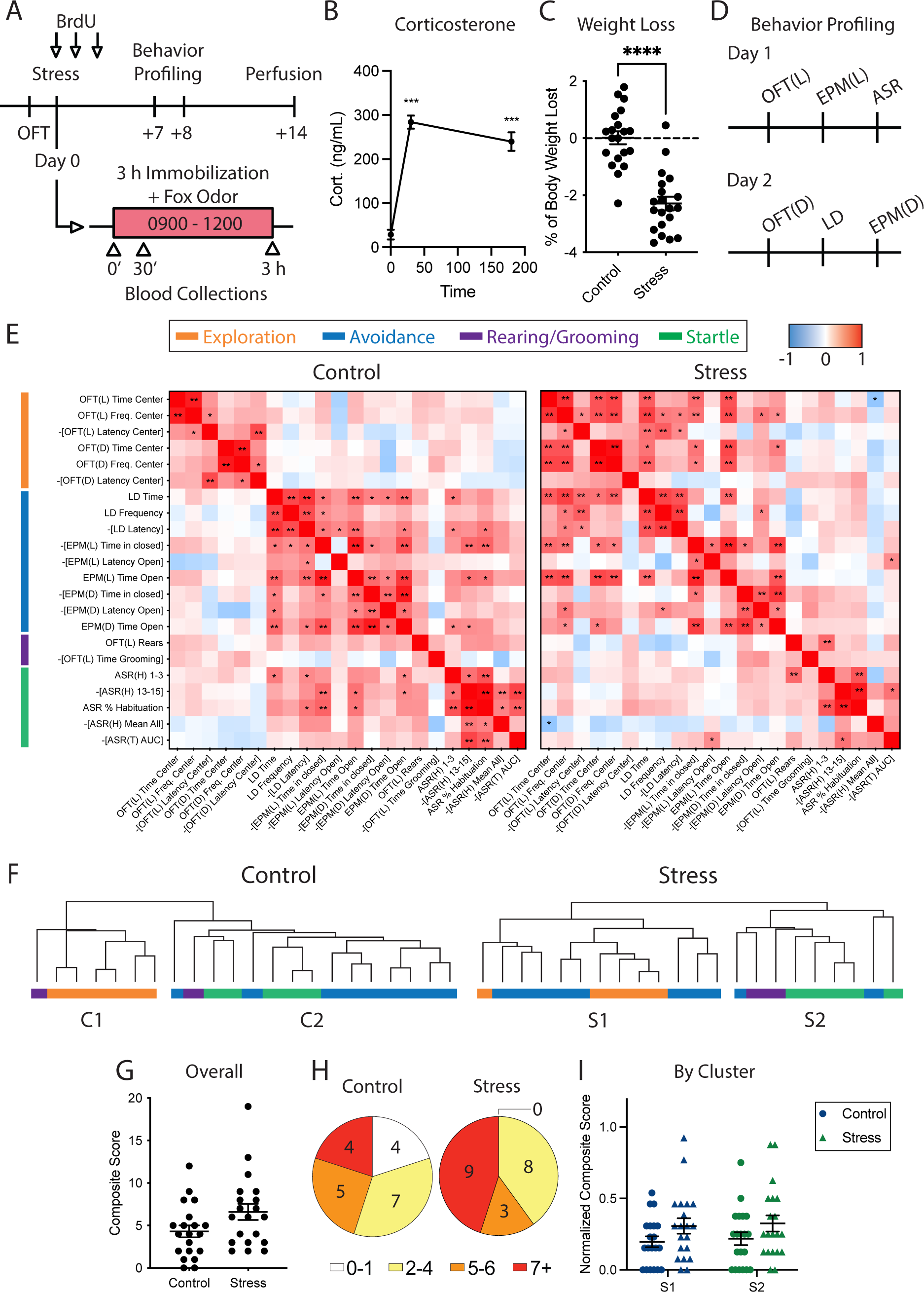
Acute, severe stress alters individual profiles of exploration, avoidance, and startle behavior in rats. **(A)** Experimental design. BrdU, bromodeoxyuridine; OFT, open field test. **(B)** Serum corticosterone at 0, 30, and 180 minutes into stress. Stress induced a 9.9-fold increase in corticosterone that persisted throughout the course of stress (mean ± SEM: time 0 = 28.60 ± 11.20 ng/mL, time 30 = 283.99 ± 15.06 ng/mL, time 180 = 239.74 ± 20.87 ng/mL; repeated measures ANOVA, F1,38 = 69.533, *p* < 0.0001). **(C)** Percent of body weight lost between the day of stress and the day after stress. Stress-exposed animals lost significantly more weight than control animals (control = 0.01 ± 0.23%, stress = -2.29 ± 0.24%; two- tailed t test, t(38) = 7.03, *p* < 0.0001). **(D)** Two-day behavior profiling design. OFT(L), open field test under bright light; OFT(D), open field test under dim light; EPM(L), elevated plus maze under bright light; EPM(D), elevated plus maze under dim light; ASR, acoustic startle response; LD, light-dark box. **(E)** Correlation matrices of behavioral measures from control and stress-exposed groups. Pearson r values are represented with red as positive and blue as negative. For ease of interpretation, latency measures, time spent in anxiolytic zones, etc. were negated to maintain a consistent directionality of valence (e.g. –[LD Latency]). Freq., frequency; ASR(H), ASR habituation phase: 1-3, average startle to stimuli 1-3; 13-15, average startle to stimuli 13-15; %Hab., percent habituation; Mean All, mean startle to all 15 110 dB stimuli. ASR(T) AUC, ASR threshold phase area under the curve. **(F)** Hierarchical clustering of behavioral measures for control and stress-exposed animals. Clusters are denoted as C1 (control 1), C2 (control 2), S1 (stress 1), and S2 (stress 2). **(G)** Composite scores of anxiety-like behavior from all tests (control: 4.3 ± 0.7, stress: 6.6 ± 1.0; two-tailed t test: t(38) = 1.94, p = 0.06; n = 20 per group). **(H)** Pie charts of animals per bin of composite scores. **(I)** Composite scores per behavioral cluster, normalized to the number of measures per cluster. \**p* < 0.05, \*\**p* < 0.01, \*\*\*\**p* < 0.0001; n = 20 for all comparisons.

**Fig. 2:**
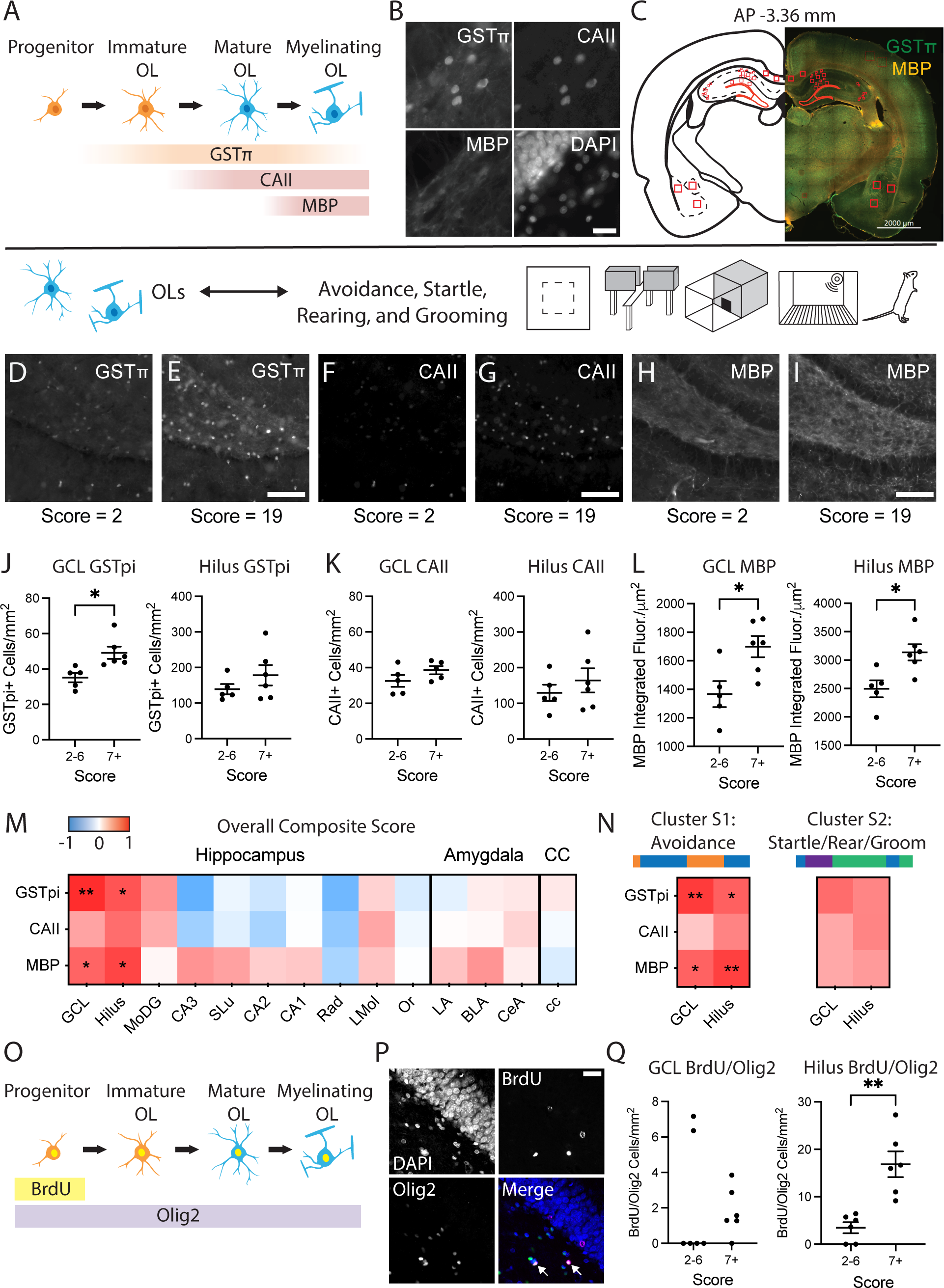
Oligodendrocytes and myelin in the dentate gyrus of the hippocampus correspond with individual outcomes of anxiety-like and avoidance behavior after acute stress. **(A)** Markers for immature and mature oligodendrocytes (OLs) and myelin selected for immunohistochemical analysis: GSTπ, glutathione S-transferase pi; CAII, carbonic anhydrase II; MBP, myelin basic protein; DAPI, 4′,6- diamidino-2-phenylindole. **(B)** Representative GSTπ, CAII, and MBP staining. Scale bar = 20 μm. **(C)** Example coronal section with hippocampal, amygdala, and corpus callosum ROIs indicated in red. Scale bar = 2000 μm. **(D-I)** Representative images of each oligodendrocyte marker from the dentate gyrus of animals with low (score = 2) or high (score = 19) composite behavior scores. Scale bar = 100 μm. **(J)** Dentate gyrus GSTπ measures from animals with low (2-6) and high (7 or more) composite behavior scores. All data are presented as mean ± SEM. Granule cell layer (GCL): low = 35.15 ± 2.64 cells/mm^2^, high = 49.16 ± 3.46 cells/mm^2^; two-tailed t test, t(9) = 3.11, *p* = 0.013. Hilus: low = 139.20 ± 14.40 cells/mm^2^, high = 178.30 ± 28.69 cells/mm^2^; two-tailed t test, t(9) = 1.14, *p* = 0.28. **(K)** CAII measures. GCL: low = 33.03 ± 3.75 cells/mm^2^, high = 38.57 ± 2.32 cells/mm^2^; two-tailed t test, t(8) = -1.26, *p* = 0.24, one high outlier removed. Hilus: low = 129.50 ± 22.95 cells/mm^2^, high = 164.40 ± 33.82 cells/mm^2^; two-tailed t test, t(9) = 0.82, *p* = 0.43. **(L)** MBP measures. GCL: low = 1366 ± 91 integrated fluorescence intensity/μm^2^, high = 1699 ± 74 integrated fluorescence intensity/μm^2^; two-tailed t test, t(9) = 2.86, *p* = 0.019. Hilus: low = 2496 ± 150 integrated fluorescence intensity/μm^2^, high = 3136 ± 142 integrated fluorescence intensity/μm^2^; two-tailed t test, t(9) = 3.09, *p* = 0.013. **(M)** Correlation matrix of composite behavior scores to oligodendrocyte and myelin measures from a subset of stress-exposed animals (Pearson correlations: n = 10-11; one high outlier removed from GCL CAII dataset; one high outlier removed from MoDG GSTπ dataset). MoDG, molecular layer of dentate gyrus; SLu, stratum lucidum; Rad, stratum radiatum; LMol, lacunosum moleculare; Or, stratum oriens; LA, lateral amygdala; BLA, basolateral amygdala; CeA, central amygdala; cc, corpus callosum. **(N)** Correlation matrices of dentate gyrus oligodendrocytes and myelin from stress-exposed animals to composite scores derived from hierarchical behavior clustering (Pearson correlations: n = 10-11; one high outlier removed from GCL CAII dataset). \**p*<0.05, \*\**p*<0.01. **(O)** Immunomarkers for quantification of oligodendrogenesis. BrdU, bromodeoxyuridine; Olig2, oligodendrocyte transcription factor 2. **(P)** Representative images of BrdU and Olig2 immunolabeling. Arrows point to colabeled cells. Scale bar = 25 μm. **(Q)** BrdU/Olig2 colabeling measures. GCL: median low = 0 cells/mm^2^, median high = 1.43 cells/mm^2^; two-tailed Mann-Whitney test, U = 14, *p* = 0.57. Hilus: low = 3.48 ± 1.16 cells/mm^2^, high = 16.86 ± 2.71 cells/mm^2^; two-tailed t test, t(10) = 4.53, *p* = 0.0011.

**Fig. 3:**
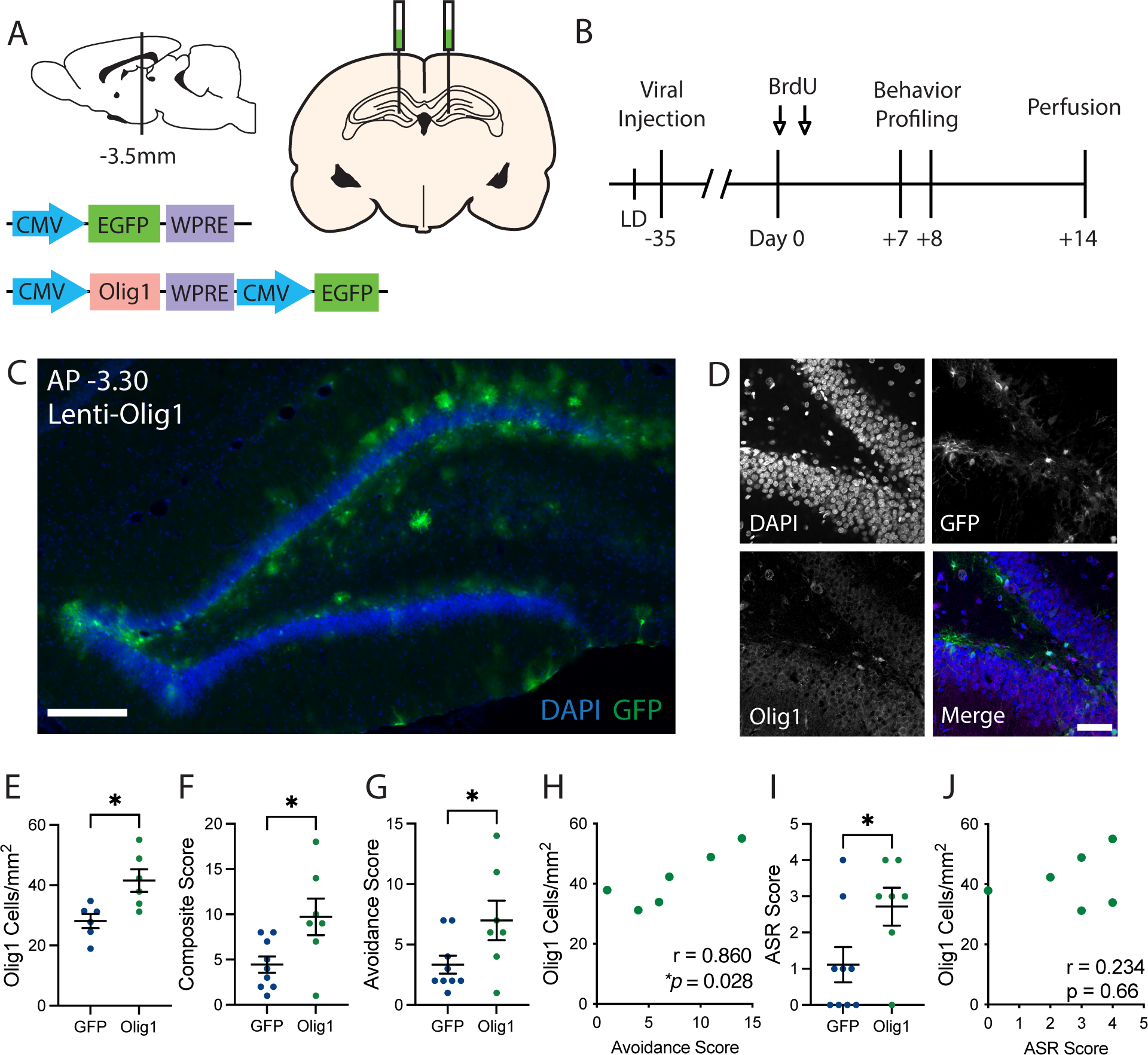
Viral upregulation of Olig1 in the dentate gyrus mimics the effects of acute stress. **(A)** Injection schematic and lentiviral constructs. CMV, human cytomegalovirus promoter; EGFP, enhanced green fluorescent protein; WPRE, Woodchuck hepatitis virus posttranscriptional regulatory element; Olig1, oligodendrocyte transcription factor 1. **(B)** Experimental timeline. BrdU, bromodeoxyuridine; LD, light- dark box. **(C)** Representative image of viral spread through the dorsal dentate gyrus. Lenti, lentivirus; DAPI, 4′,6-diamidino-2-phenylindole; GFP, green fluorescent protein. Scale bar = 200 μm. **(D)** Representative images of GFP and Olig1 immunohistochemistry 7 weeks post-injection. Scale bar = 50 μm. **(E)** Dentate gyrus Olig1 cell densities (GFP: 28.11 ± 2.33 cells/mm^2^, Olig1: 41.54 ± 3.72 cells/mm^2^; two-tailed t test: t(10) = 3.06, *p* = 0.012; n = 6 per group). **(F)** Composite behavior scores (GFP: 4.4 ± 0.9, Olig1: 9.7 ± 2.0; two-tailed t test: t(14) = 2.58, *p* = 0.022; nGFP = 9, nOlig1 = 7). **(G)** Composite behavior scores from avoidance tests (GFP: 3.3 ± 0.7, Olig1: 7.0 ± 1.6; two-tailed t test: t(14) = 2.21, *p* = 0.044; nGFP = 9, nOlig1 = 7). **(H)** Pearson correlation of Olig1 cell densities to avoidance composite scores from Lenti-Olig1 animals (Pearson correlation: r = 0.86, *p* = 0.028; n = 6). **(I)** Composite behavior scores from the acoustic startle response (ASR) test (GFP: 1.1 ± 0.5, Olig1: 2.7 ± 0.5; two-tailed t test: t2,14 = 2.24, *p* = 0.042; nGFP = 9, nOlig1 = 7). **(J)** Pearson correlation of Olig1 cell densities to composite startle scores from Lenti-Olig1 animals (Pearson correlation: r = 0.23, *p* = 0.66; n = 6). For all plots, individual data points are shown with mean ± SEM, \**p* < 0.05.

### Open field test (OFT)

Each rat was placed in an unenclosed plastic box (50 w x 50 l x 58 h in cm) and was given 10 minutes to explore the arena. All animals were placed along a wall at the beginning of the test. Behavior was recorded with cameras positioned above the arena and connected to GeoVision software (GeoVision Inc., Taiwan). Behavior was scored as latency to, frequency, and total amount of time spent in the center of the box (designated by a 15 x 15 cm square) EthoVision software (Noldus, Leesburg, VA). In addition, we quantified the duration of rearing (upright posture with both forepaws lifted off the ground) and self-grooming from the OFT Light. The OFT Light was conducted under full lighting (280 lux). The OFT Dim was conducted in a different but identically structured box in the same room under 15 lux. Arenas were cleaned with 1% acetic acid followed by Formula 409 All Purpose Cleaner after each animal.

### Elevated plus maze (EPM)

Rats were allowed to explore an EPM for 10 min (arm dimensions: 10 w x 60 l in cm; closed arm enclosed by walls 51 cm in height; apparatus elevated 50 cm off the ground). Behavior was recorded by a JVC Everio camera (JVCKENWOOD, Tokyo, Japan) mounted above the apparatus. The criterion for open arm exploration was placement of more than half of the body (and both forepaws) into the open arm. Latency to and total time spent in the exposed open arms, as well as time spent in the protected closed arms, were quantified by observers blind to condition. The EPM Light was conducted at 240 lux, while the EPM Dim was conducted at 125 lux. The apparatus was cleaned with 1% Process NPD Disinfectant (STERIS Life Sciences) after each animal.

### Light-dark box (LD)

Each rat was placed in a structure consisting of an enclosed dark box separated by a divider with a small door leading to an unenclosed light box (each box 15 w x 15 l x 8 h in inches). All animals were placed into the dark half of the box and given 10 min to explore. Behavior was recorded by a JVC Everio camera (JVCKENWOOD, Tokyo, Japan) mounted above the apparatus. Measures for latency to, frequency, and total time spent in the exposed side were quantified via EthoVision software or observers blind to condition. The arena was cleaned with 70% ethanol after each animal.

### Acoustic startle response (ASR)

Each rat was placed into an isolated Coulbourn sound-attenuating fear conditioning chamber (12 w x 10 l x 12 h in inches) and was exposed to 5 min of background noise (∼55 dB). This was followed by 70-110 dB white noise pulses lasting 10 ms, with an inter-stimulus interval of 15-30 s. All tones were calibrated with a handheld decibel meter each day prior to testing. Behavior was recorded over two different trials: habituation (110 dB tones presented 15 times to assess initial responses and subsequent habituation) and threshold determination (70-110 dB tones presented in pseudo-random order, with each tone played 5 times in total). Behavior was recorded using a Coulbourn Instruments camera connected to a computer with FreezeFrame software (Coulbourn Instruments, Whitehall, PA). The boxes were cleaned with 70% ethanol after each animal. Startle behavior was assessed by EthoVision software analysis of activity change (measured as percent pixel change from frame to frame). The amplitude of startle was quantified as the maximum activity minus baseline activity in the 50 ms surrounding the startle pulse. From the habituation phase, we quantified initial startle (mean startle amplitude to stimuli 1-3); final startle (mean startle amplitude to stimuli 13-15); mean startle (mean of all startle amplitude scores across all 15 stimuli); and sensitization, calculated as 100*(final startle - initial startle)/(initial startle). Habituation is, therefore, Sensitization*(-1). From the threshold phase, we averaged startle scores from the 5 presentations of each of the decibel levels (70, 80, 90, 100, and 110 dB) to generate a startle threshold curve for each animal, and we quantified the area under the curve (AUC).

### Composite behavior scoring

To standardize and quantify behavior across multiple behavior tests, we adapted the method of Cutoff Behavioral Criteria developed by Cohen and Zohar (49). For each measure, a behavioral cutoff criterion was defined as the 20^th^ percentile of the control distribution. For measures in which greater scores indicate greater anxiety-like behavior (latency to an anxiogenic zone, time spent in an anxiolytic zone), the 80^th^ percentile of the control group was used. Binary scoring was applied: Animals falling outside the criterion were marked as “affected” and received a score of 1 for that measure. Scores were then summed across all tests. High scores represent consistent anxiety-like behavior across all tests. We compared these scores to 3 other methods of composite scoring: Z scoring, principal component analysis (PCA), and ranks. For Z scoring, we converted all measures to Z scores using the mean and standard deviation of the control group; Z scores were then averaged across all measures from all tests. For PCA, we conducted a PCA on all measures from all animals via custom Python scripts and recorded each animal’s value from the first principal component. For ranks, for each measure, we sorted all animals in ascending order and assigned each animal a rank for that measure from 0 (least anxiety-like behavior) to 40 (highest anxiety-like behavior). For example, the animal with the least time spent in the center of the OFT received a rank of 40. Ranks were then averaged across all measures from all tests. Each of these composite scoring methods (Z scores, first principal component values, and average ranks) were compared to composite cutoff scoring via Pearson correlations.

### Fear conditioning cohort

A separate cohort of 12 male rats was used to test repeated avoidance profiling coupled with fear conditioning. All animals underwent a one-day profiling battery consisting of the OFT Light, EPM Light, and LD with 10 minutes in the home cage in between each test (day -19 relative to stress). Eight days later (day -11), the animals went through a second one-day behavior profiling battery. Rats went through the acute, severe stress paradigm (day 0) with blood sampling 11 days later, followed by a third avoidance profiling battery 7 days later (day +7). On day +8 post-stress, all animals underwent a 3-day fear conditioning protocol. On the first day, animals were placed in a Coulbourn sound-attenuating fear conditioning chamber (12 w x 10 l x 12 h in inches) with an electrified grid floor. Animals were allowed 5 min to acclimate to the box. Following acclimation, 10 unsignaled, 1 mA, 1 s duration shocks were delivered with an inter-stimulus interval of 15-120 s. Rats were left in the chamber for 3 min after the last shock and then returned to the home cage. On the second day (day +9), animals were placed back into the fear context without shock and underwent 5 extinction trials lasting 10 min each. Inter-extinction trial intervals were ∼90 min. On the third day (day +10), animals were again placed in the fear context without shock for a single 10 min extinction-retention test (Probe). The fear chamber was cleaned with 70% ethanol in between each animal. The time spent freezing in each trial was quantified by observers blind to condition. Extinction was quantified as area under the curve (AUC) from the 5 extinctions trials. Animals underwent the ASR test on day +11, and on day +14, animals underwent a final 15 min LD test prior to perfusion 15 min later.

### Olig1 lentivirus and stereotaxic injections

To test whether hippocampal oligodendrogenesis contributes to avoidance behavior, two 3^rd^ generation lentiviral expression vectors were designed with VectorBuilder (Santa Clara, CA). The control vector was designed to express enhanced green fluorescent protein (EGFP) under the human cytomegalovirus (CMV) promoter. The experimental vector was a bicistronic design with the CMV promoter driving expression of mouse Olig1 (NM_016968.4) followed by a second CMV promoter driving EGFP. Plasmids were packaged by the UC Berkeley High Throughput Screening Facility in HEK293T cells. Cells were transfected via jetPRIME (VWR, Radnor, PA) with the viral plasmid and 3 helper plasmids (pCMV-VSVG, pMDL, and pRSV-Rev). The resulting virus-containing media was collected 48 h and 72 h after transfection. Virus was precipitated with 5% PEG and 0.15M NaCl and centrifuged at 3000 g for 15 min at 4°C. The precipitated viral pellet was resuspended in ice cold phosphate-buffered saline (PBS), aliquoted, and stored at -80°C. Viral titer was determined with an ABM qPCR Lentivirus Titration Kit (ABM, Vancouver, B.C., Canada). Titers were 10^6^-10^7^ infectious particles/mL.

For viral injections into the dentate gyrus, animals were injected with 0.05 mg/kg buprenorphine and anesthetized with isoflurane (2-3%). Using a stereotaxic frame, bilateral craniotomies were made at the following coordinates: -3.5 mm anterior/posterior, +/-2.2 mm medial/lateral relative to bregma, and -3.3 mm relative to dura. For viral infusion, 1.3 µL of virus was infused at a rate of 0.1 µL/min into each hemisphere. After the injection, the needle was left in place for five minutes to allow for viral diffusion, then was slowly removed to minimize viral infection along the needle tract. One animal from the Lenti- Olig1 group was removed from further analysis due to unilateral injection failure. One animal from the Lenti-GFP control group was removed for post-surgical complications (chronic swelling around the wound site).

### Perfusion

Rats were deeply anesthetized with Euthasol Euthanasia Solution (Virbac AH, Inc.) and transcardially perfused with 0.9% saline followed by ice-cold 4% paraformaldehyde (PFA) in 0.1 M phosphate-buffered saline (PBS). Brains were post-fixed for 24 hours at 4°C in 4% PFA and equilibrated in 30% sucrose in 0.1 M PBS. They were then stored at -80°C. The brains were sliced into free-floating 40 µm sections on a cryostat in a 1 in 12 series and were subsequently stored at -20°C in antifreeze solution.

### Immunohistochemistry and fluorescence microscopy

Immunohistochemical staining was conducted on free-floating sections. All antibodies were tested and validated by including a negative control (no primary antibody applied) during staining. On the first day, slices were washed in tris-buffered saline (TBS) and blocked with 3% normal donkey serum (NDS) in TBS with 0.3% Triton-X100 for one hour at room temperature. The slices were then incubated overnight at 4°C in primary antibodies diluted in TBS with 0.3% Triton-X100 and 1% NDS. Primary antibody incubation for Olig1 was extended to 72 hours at 4°C. Primary antibodies were rat anti-myelin basic protein (MBP; 1:500; Abcam, Cambridge, UK), rabbit anti-glutathione S-transferase π (GSTπ; 1:1000; MBL International, Woburn, MA), sheep anti-carbonic anhydrase II (CAII; 1:10000; AbD Serotec, Hercules, CA), rabbit anti-Olig1 (1:1000; MilliporeSigma, Burlington, MA), and chicken anti-green fluorescent protein (GFP; 1:750; Abcam, Cambridge, UK). On the second day of staining, the slices were rinsed in TBS and incubated in 1:500 diluted fluorophore-conjugated secondary antibodies for 2 hours at room temperature in blocking solution with 1% NDS. Secondary antibodies were Alexa Fluor 647 donkey anti-sheep, Cy3 donkey anti-rat, Alexa Fluor 488 donkey anti-rabbit, Alexa Fluor 594 donkey anti-rabbit, and Alexa Fluor 488 donkey anti-chicken (all from Jackson ImmunoResearch, West Grove, PA). Lastly, the sections were rinsed in TBS and stained with the nuclear marker DAPI (1:40,000 in TBS). Stained sections were mounted onto glass slides and coverslipped with DABCO antifading medium.

For BrdU staining, the first day of staining proceeded as detailed above, and slices were incubated with rabbit anti-Olig2 (1:1000; MilliporeSigma, Burlington, MA) overnight at 4°C in blocking solution with 1% NDS. On the second day of staining, the slices were rinsed in TBS and incubated in 1:500 Alexa Fluor 488 donkey anti-rabbit for 2 hours at room temperature in blocking solution with 1% NDS. Sections were then rinsed in TBS and incubated in 4% PFA for 10 min. Sections were rinsed in TBS 3 times, rinsed in 0.9% saline once, and then incubated in 2N HCl at 37°C for 30 minutes. Sections were then rinsed in TBS and incubated in blocking solution with 3% NDS for 1 hour at room temperature. Sections were then incubated overnight at 4°C in blocking solution with 1% NDS and primary rat anti-BrdU (1:500, Abcam, Cambridge, UK). On the third day, sections were rinsed and incubated in blocking solution with secondary Cy3 donkey anti-rat (Jackson ImmunoResearch, West Grove, PA) for 2 hours at room temperature. Lastly, the sections were rinsed in TBS and stained with the nuclear marker DAPI (1:40,000 in TBS). Stained sections were mounted onto glass slides and coverslipped with DABCO antifading medium.

Imaging of slides was conducted under a 40x objective using a Zeiss 510 AxioImager microscope (Zeiss, Oberkochen, Germany) with MetaMorph software (Molecular Devices, San Jose, CA) and a 10x air objective on a Zeiss AxioScan Slide Scanner (AxioScan.Z1, Zeiss, Oberkochen, Germany) with Zen software (Zeiss). Regions of interest were scanned and automatically stitched. For the dentate gyrus, sections from Bregma -2.9 to -5.28 were imaged, and the granule cell layer (GCL) and hilus (defined as the region within the GCL blades and medial to the stratum pyramidale of CA3) were traced. Cells were counted from 4-8 hemispheres of the dorsal dentate gyrus from each animal with Fiji software (50). Overall DG measures were collected by summing measures from the hilus and GCL. For other brain regions, ROIs of standardized size (75 x 75 µm to 300 x 300 µm) were collected from both hemispheres of 2-6 sections. Cell bodies were counted via automated thresholding and particle analysis in Fiji, and output was visually confirmed by observers blind to condition. This allowed for high throughput cell counting from many regions. For all regions, myelin content was measured from the raw images by the integrated fluorescence intensity of MBP immunofluorescence. All measures for all regions were averaged across sections and normalized to area in mm^2^ (cell densities) or µm^2^ (MBP intensity).

### Study design - Humans

#### MRI Data Acquisition and Processing

Our original publication in humans, Chao et al., 2015 (46), includes detailed methods regarding sample accrual, diagnostic classification, brain structural data acquisition, and analyses. Briefly, 38 male veterans (19 PTSD+, 19 PTSD-) were studied at the Center for Imaging of Neurodegenerative Diseases at the San Francisco Veterans Affairs Medical Center. Screening exclusions included a history of psychosis or bipolar disorder, alcohol abuse/dependence within the last 12 months, and substance abuse/dependence within the last six months. PTSD diagnoses and symptoms were obtained using the Clinician Administered PTSD Scale (CAPS) by trained clinicians. Structural magnetic resonance imaging (MRI) scans were acquired using a Bruker/Siemens Med-Spec 4T MRI system (Bruker BioSpin, Ettlingen, Germany). The image series acquired included a T1-weighted magnetization prepared gradient echo sequence and a volumetric T2-weighted turbospin echo sequence. The images were processed and assessed for myelination levels using the methods developed by Glasser and Van Essen, 2011 (51), validated in 1555 brains by Shafee, Buckner, and Fischl, 2015 (52), and outlined by Chao et al., 2015 (46). The FreeSurfer version 4.5 tissue boundary delineations were used to assign regions of interest for the hippocampus, amygdala, and corpus callosum (53).

#### Statistical Analysis

All data are presented as mean ± standard error of the mean (SEM). Analyses were performed using IBM SPSS 19 (SPSS, Inc., Chicago, IL) and GraphPad Prism version 9.1.1 for Mac OS X (GraphPad Software, San Diego, California USA, www.graphpad.com). Hierarchical clustering was performed with the Seaborn package in Python (Version 3.7.1, Python Software Foundation, Wilmington, DE). Two animals were excluded from the virus cohort for surgical complications or unsuccessful viral targeting (detailed above); otherwise, all animals were included. For correlations between brain measures and behavior scores, we screened datasets for outliers using the ROUT method in Prism with Q-value set to 1% (54). Instances in which outliers are excluded are noted in figure legends and the supplemental Excel file. Two sample comparisons were performed by two-sided Student’s t-test, and multiple group comparisons were conducted by ANOVA, followed by Tukey’s post-hoc testing to compare individual groups when a main effect was detected. Greenhouse-Geisser corrections were used when the assumption of sphericity was broken, as indicated by Mauchly’s test of sphericity. To compare the relationships between measures, we conducted Pearson correlations. In all tests, the alpha value was set at 0.05. For human work, descriptive statistics were performed on the demographic and clinical characteristics across the groups using both the Student’s t-test for continuous variables and the Fisher’s exact test of independence for categorical variables. The relationship between PTSD symptom severity (i.e., CAPS) and hippocampal myelin content was examined with Spearman’s Rank-Order correlation. Descriptive and hypothesis-testing statistics can be found in the supplemental Excel file.

## Results

### Exposure to acute, severe stress yields a spectrum of anxiety-like behavior in adult rats

To model exposure to acute, severe trauma, we subjected adult, male Sprague-Dawley rats to immobilization with predator scent (fox urine) (Fig. 1A). This is an established model with ethological validity that elicits a large corticosterone response in the rat (28,55–58). Twenty rats underwent stress, while twenty served as controls and remained minimally disturbed. Animals exposed to stress had a dramatic increase in circulating corticosterone levels by 30 minutes and 3 hours into exposure (Fig. 1B) and lost significantly more weight than control animals by the day after stress (Fig. 1C), confirming that this paradigm was effective at eliciting an acute, physiological stress response.

Following acute trauma exposure in rodents and humans, the majority of individuals gradually recover to baseline levels of fear and anxiety, while a subset of susceptible individuals displays persistent changes to behavior (4, 49). To identify stress-susceptible rats, we allowed animals to remain in the home cage with minimal disturbance for 7 days after stress. We then characterized behavioral changes with a suite of tests spanning the domains of exploration, avoidance, startle, rearing, and grooming (Fig. 1D). This paradigm identifies sub-populations of stress-exposed animals displaying persistent and stable changes to different anxiety-like behaviors (49,59,60). Assays included the open field test (OFT) under highly anxiogenic (brightly lit: OFT Light) and low anxiogenic (dimly lit: OFT Dim) conditions, elevated plus maze (EPM) under bright and dim conditions, light-dark box test (LD), and acoustic startle response (ASR) test. For OFT, EPM, and LD tests, we quantified frequency, duration, and latency of visits to anxiogenic and anxiolytic zones. From the OFT Light, we also quantified rearing and grooming behavior, both of which have been used as ethological measures of general anxiety-like behavior in the rat (61, 62). From the ASR test, we quantified habituation (and, conversely, sensitization) to repeated startle stimuli, mean startle response, and the startle response curve to tones of increasing decibels (48).

We first sought to characterize this multi-dimensional behavioral data set. Broadly, animals exhibited considerable variability in each individual metric (Fig. S1). To determine how behavioral metrics from separate domains relate to each other under baseline conditions and following stress exposure, we correlated behavioral measures within and across assays (Fig. 1E). Interestingly, in the control group, avoidance assays showed several cross-assay correlations to startle measures, but not to exploration measures. These correlations were confirmed by unbiased hierarchical clustering (Fig. 1F, Fig. S2). In the control group, the first order branching yielded 2 clusters. Control Cluster 1 consisted of all OFT measures and grooming, while Control Cluster 2 consisted of all avoidance and startle measures, as well as rearing. Conversely, correlations and hierarchical cluster membership were altered in stress-exposed animals when compared to controls. Specifically, avoidance assays showed cross-assay correlations to exploration, but not to startle measures. Stress Cluster 1 (S1) consisted of all OFT measures and nearly all avoidance measures, while Stress Cluster 2 (S2) consisted of all startle measures, as well as rearing, grooming, and 2 avoidance latency measures. Broadly, the rodent-specific anxiety-like measures of rearing and grooming showed no strong relationships to measures from other domains in either control or stress-exposed groups. Together, this reveals that acute stress exposure shifts behavioral patterns such that exploration of the open field aligns with measures of avoidance, but not startle, in stress-exposed animals.

Clinical scores such as the Clinician Administered PTSD Scale (CAPS) provide a numeric value that encapsulates many individual, yet related, measures, providing greater descriptive power than a single measure in isolation. To generate a summative index for behavior across all tests, we adapted the concept of Cutoff Behavioral Criteria to assign a binary score for each anxiety-like behavioral measure (Fig. S3) (48, 49). Briefly, quantitative measures were compared to the control group distribution and given a score of 0 or 1 depending on whether they fell below a cutoff criterion, defined here as the 20th percentile of the control group (Fig. S1 and Fig. S3A-B). This cutoff was chosen to capture animals with the most extreme behavioral responses and to model the prevalence rates of high anxiety behavior seen in rats and clinical populations (63). Binary outcomes were then summed across all measures (Fig. S3C). High composite scores reflect that an animal consistently displayed anxiety-like behavior across many measures when compared to control animals. Similar methods have been used to classify rats subjected to predator exposure, and a recent study used cutoff scoring to classify distinct behavioral phenotypes in mice subjected to chronic unpredictable stress (64, 65). Here, we found that cutoff-based composite scores were strongly correlated with values from 3 separate methods of quantifying individual variability: Z scoring, principal component analysis, and averaged ranks across measures, providing further validity for this technique (Fig. S3D-F). Using this method, we found that stress-exposed animals displayed a wide range of composite scores that trended higher than controls (p = 0.06, Fig. 1G). Interestingly, scores from stress-exposed animals spanned a large range from minimally affected (composite score = 2) to highly affected (composite score = 19). We then binned scores according to the mean and standard deviation of control scores (4.3 ± 3.2) to yield four bins with cutoffs of 1, 4, and 7 (bins: 0-1, 2-4, 5-6, and 7+). Notably, no stress-exposed animals fell into the lowest bin of scores (0–1), while the percentage of stress-exposed animals falling into the highest bin (7+) was more than doubled compared to controls (Fig. 1H). To determine whether the upward shift in scores was driven by a particular behavioral cluster, we generated composite scores using measures from either cluster S1 or S2. We found that stress-exposed animals exhibited upward trends in composite scores for both clusters (Fig 1I). Altogether, this suggests that rats display individual variability in both the type and severity of outcomes after exposure to acute severe stress, with specific shifts in the behavioral domains of avoidance and startle.

### Oligodendrocytes and myelin in the dentate gyrus correspond to individual avoidance outcomes in stress- exposed animals

Next, we tested whether oligodendrocyte cell density and myelin content in select white and gray matter regions of interest corresponded to the behavioral variation of individual anxiety-like outcomes after stress. In a subset of stress-exposed animals (n = 11) chosen to span the full range of composite scores, we analyzed cellular immuno-markers of the oligodendrocyte lineage, including glutathione S-transferase π (GSTπ, an intermediate-to-late-stage oligodendrocyte marker), carbonic anhydrase II (CAII, a marker of mature oligodendrocytes), and myelin basic protein (MBP, a marker of myelination) (Fig. 2A,B). We quantified the number of oligodendrocyte cell bodies and myelin staining intensity in multiple hippocampal subregions, including the granule cell layer (GCL) and hilus (polymorphic layer) of the dentate gyrus, molecular layer, CA3, CA2, CA1, stratum lucidum, stratum radiatum, stratum lacunosum moleculare, and stratum oriens. We further quantified oligodendrocyte density and myelin staining in the amygdala lateral subdivision (LA), basolateral subdivision (BLA), and central subdivision (CeA), as well as in the brain’s major white matter track, the corpus callosum (Fig. 2C).

Interestingly, greater oligodendrocyte cell density and greater myelin amounts specifically in the dentate gyrus of the hippocampus corresponded to greater anxiety-like behavior after stress (Fig. 2D-I). Animals classified as having the highest levels of anxiety-like behavior (composite scores of 7+) had significantly greater GSTπ cell density in the GCL, but not the hilus, when compared to animals with low scores (scores of 2-6; Fig. 2J). In contrast, oligodendrocyte CAII measures were not significantly different across group outcomes (Fig. 2K). Animals with high composite scores also had significantly greater MBP fluorescence in both the GCL and the hilus (Fig. 2L). Remarkably, a correlation matrix revealed that composite scores in stress-exposed animals strongly correlated with GSTπ and MBP in the GCL and hilus (Fig. 2M), suggesting that oligodendrocytes and myelin in the dentate gyrus not only correspond to group-level differences in anxiety-like behavior, but also serve as correlative biomarkers of individual variability after stress. This was not the case for any other region of the hippocampus, amygdala, or corpus callosum, nor was it the case for control animals (Fig. S4). We tested whether composite scores derived from either cluster S1 (Avoidance) or S2 (Startle, Rearing, Grooming) drive the relationship between anxiety-like behavior and dentate gyrus oligodendrocytes. In stress-exposed animals, we found that dentate gyrus GSTπ and MBP significantly positively correlated with composite scores from cluster S1, but not cluster S2, suggesting that dentate gyrus oligodendrocytes and myelin correspond most strongly with individual outcomes of avoidance behavior (Fig. 2N). Overall, these experiments suggest that oligodendrocyte and myelin markers in the rat dentate gyrus correspond to stress-induced anxiety-like outcomes following exposure to a single, severe stressor.

We have previously demonstrated that stress and glucocorticoids can induce oligodendrogenesis from neural stem cells in the dentate gyrus (32). Here, we tested whether oligodendrocytes born in the timeframe surrounding the stressor corresponded to individual outcomes of anxiety-like behavior. We injected the thymidine analog BrdU immediately after stress as well as on the 2 days following stress to label actively dividing cells. After perfusion, we stained for BrdU and the oligodendrocyte transcription factor Olig2 (Fig. 2O,P). Animals with high composite scores had significantly greater BrdU/Olig2 cell densities in the hilus than animals with lower scores (Fig. 2Q). The correlation between BrdU/Olig2 cells and composite behavior scores was trending, but not statistically significant with our sample size of 11 (r = 0.522, p = 0.08). Together, this may suggest that oligodendrogenesis and/or oligodendrocyte survival in the dentate gyrus correspond with higher anxiety-like behavior after acute, severe stress.

### Manipulation of hippocampal oligodendrogenesis mimics stress exposure

Given the group-level and correlative relationships between dentate gyrus oligodendrocytes and behavior, we next tested whether manipulating hippocampal oligodendrocytes is sufficient to induce changes in anxiety-like behavior. We developed a lentiviral construct to induce ectopic overexpression of the oligodendrogenic transcription factor Olig1 (*Lenti-CMV-mOlig1-CMV-EGFP*) or GFP as a control (*Lenti-CMV-EGFP*; Fig. 3A). We stereotactically delivered virus to the dorsal dentate gyrus of male rats (nGFP = 9, nOlig1 = 7; Fig. 3A). After six weeks of recovery to allow for viral expression, animals underwent two-days of behavior profiling (Fig.3B).

Viral delivery and expression in the dentate gyrus were confirmed by GFP expression (Fig. 3C-D). We found that viral expression in Lenti-Olig1 animals was largely restricted to the subgranular zone and molecular layer, with some additional expression in the hilus (Fig. 3C-D). To confirm effective upregulation of Olig1 expression, we quantified Olig1 cell density in the GCL and hilus in a subset of animals. Rats that received Lenti-Olig1 virus indeed showed significantly more Olig1 cell density in the dentate gyrus compared to Lenti-GFP animals (Fig. 3E). We calculated composite scores of anxiety-like behavior using Lenti-GFP animal behavior as the reference control distribution. We found that animals that received Lenti- Olig1 virus displayed significantly enhanced anxiety-like behavior scores when compared to Lenti-GFP animals (Fig. 3F and Fig. S5). To determine which behavioral domains were altered by viral delivery, we generated composite subscores for avoidance measures and startle measures separately. Lenti-Olig1 animals displayed significantly greater composite avoidance scores than Lenti-GFP animals (Fig. 3G), and these scores were significantly positively correlated with Olig1 cell density (Fig. 3H), echoing the findings from our first experiment. Lenti-Olig1 animals also displayed greater composite startle scores relative to Lenti-GFP animals (Fig. 3I); however, these scores were not correlated with Olig1 cell densities (Fig. 3J). Together, these findings suggest that Olig1 ectopic overexpression in the dorsal dentate gyrus increases both avoidance and startle behavior in rats. Hence, manipulation of the oligodendrocyte population in the absence of exposure to stress was sufficient to phenocopy stress-induced place avoidance and to enhance startle. These findings highlight the relationship between hippocampal oligodendrogenesis and behavioral outcomes and suggest that oligodendrogenesis can contribute to the pathogenesis of enhanced stress reactivity.

### Increased myelin content in the amygdala and hippocampus is correlated with enhanced contextual fear learning

In addition to avoidance behavior and hypervigilance, trauma can induce long-lasting changes in fear learning and fear extinction (66, 67). In our first experiments, we identified region-specific relationships between oligodendrocyte and myelin content and the behavioral domains of avoidance and startle. Thus, we next sought to expand our behavioral analyses to include fear learning and to test whether regional oligodendrocytes and myelin also correspond to individual outcomes of fear following exposure to acute, severe stress. Furthermore, we hypothesized that baseline behavioral profiles may predict an individual’s susceptibility to stress. Thus, we sought to test whether avoidance behavior prior to stress could predict individual outcomes in fear behavior following stress exposure.

To test these hypotheses, we conducted a study in which 12 male rats underwent behavioral profiling (OFT + EPM + LD) at 3 time points -- twice before stress (t1 = -19 and t2 = -11 days) and once following stress (t3 = +7 days) (Fig. 4A). To probe for individual variation in fear learning following stress, animals underwent fear conditioning on day +8 after stress. We quantified freezing behavior during the initial shock session. In addition, contextual fear memory and fear extinction were quantified by measuring freezing behavior during 5 extinction trials, and extinction memory was quantified by measuring freezing during an extinction probe trial. We found that animals displayed a wide range of time spent freezing in each of the trials, suggesting that rats display individual variability in the severity of fear expression and fear learning outcomes after acute, severe stress (Fig. 4B).

**Fig. 4:**
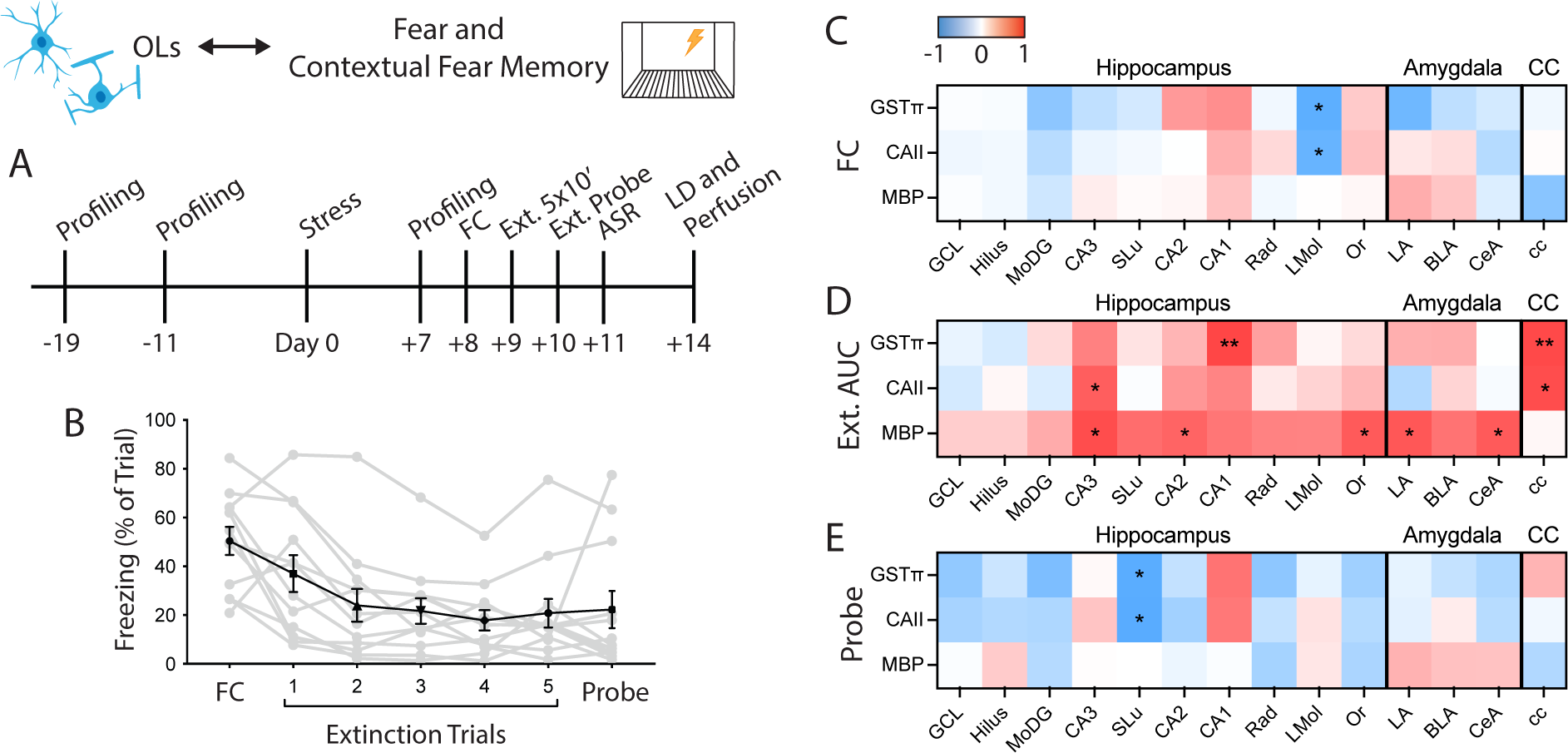
Amygdala myelin is positively correlated with contextual fear memory after acute stress in rats. **(A)** Experimental design. **(B)** Freezing across fear conditioning, extinction, and extinction probe trials as percent of total trial time. Individual animals are shown in gray with mean ± SEM freezing of all 12 animals shown in black. FC, Fear conditioning. **(C)** Correlation matrix of freezing during fear conditioning (FC) trial to oligodendrocyte and myelin measures (Pearson correlations: n = 12). **(D)** Correlation matrix of area under the curve (AUC) of extinction freezing traces to oligodendrocyte and myelin measures (Pearson correlations: n = 12). **(E)** Correlation matrix of freezing during extinction probe trial to oligodendrocyte and myelin measures (Pearson correlations: n = 10; two high outliers removed from probe freezing dataset). \**p* < 0.05, \*\**p* < 0.01.

We then asked whether individual profiles of avoidance behavior prior to and following stress predict fear outcomes. Broadly, we found that fear conditioning measures did not correlate with avoidance measures from either before or after stress (Fig. S6A-B). In addition, we compared fear conditioning measures to startle behavior after stress and again found that the majority of measures were not significantly correlated (Fig. S6C). Altogether, these findings suggest that avoidance, startle, and fear are separate domains of behavior in this model of acute, severe stress and that an individual’s outcome in one behavioral domain does not predict its outcome in another domain.

To test whether oligodendrocytes and myelin correspond to individual differences in fear-like behavior, we quantified GSTπ and CAII cell density and MBP fluorescence intensity from white and gray matter regions of interest as described above and correlated these measures to individual fear-like behavior metrics. This revealed that, in contrast to avoidance and startle behavior, oligodendrocytes and myelin in the dentate gyrus of the hippocampus showed no significant relationships to fear conditioning or extinction measures (Fig. 4C-E). Rather, MBP immunofluorescence in the CA3, CA2, and stratum oriens subregions of the hippocampus were positively correlated with freezing during extinction trials (Fig. 4D). In addition to these hippocampal regions, MBP immunofluorescence in the LA and CeA of the amygdala positively correlated with freezing behavior during extinction trials, but not to the initial fear conditioning trial or extinction probe (Fig. 4C-E). These findings suggest a potential role for hippocampal and amygdala myelin levels in contextual fear memory following exposure to a single, severe stressor in rats.

### Hippocampal and amygdala myelin correspond to avoidance and fear in trauma-exposed male veterans

Previously, we showed that hippocampal myelin is positively correlated with overall PTSD symptom severity as measured by CAPS scores (46). CAPS IV scores encompass symptoms for avoidance (Avoidance sub-score), fear (Re-experiencing sub-score), and hypervigilance (Hyperarousal sub-score) (68). Thus, following our findings in the rat hippocampus and amygdala, we hypothesized that estimates of myelin content in these same regions would regionally correspond to individual behavioral outcomes in human veterans exposed to trauma. To test this, we extended our previous work and correlated myelin estimates from the hippocampus, amygdala, and corpus callosum with individual sub-components of CAPS scores.

As previously shown (46), there is a significant, positive correlation between overall CAPS scores and myelin estimates in the hippocampus (Table 1). Here, we additionally found that myelin estimates from the amygdala, but not the corpus callosum, correlated with overall CAPS scores. When further comparing myelin estimates to categorical sub-scores, we found that both hippocampal and amygdala myelin, but not corpus callosum myelin, were positively correlated with Avoidance and Hyperarousal sub-scores, but not Re-experiencing sub-scores.

**Table 1:**
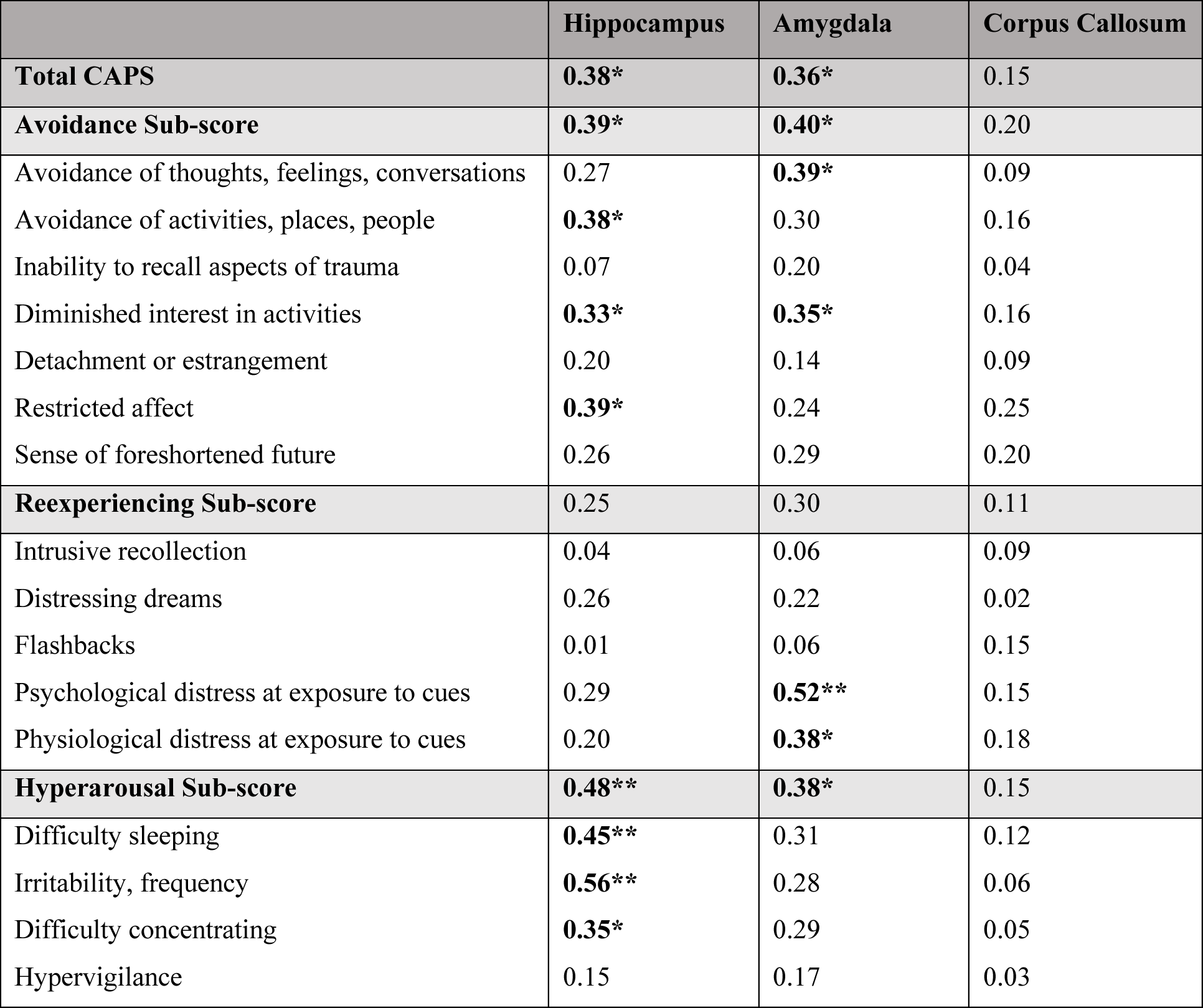
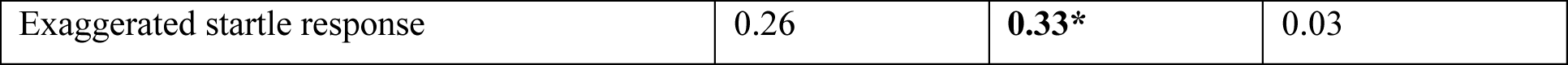
Correlations of human regional myelin estimates to sub-components of the Clinician Administered PTSD Scale (CAPS). Myelin estimates were measured via T1/T2-weighted imaging from 38 trauma-exposed male veterans, as described in Chao *et al.*, 2015. 19 trauma-exposed veterans were diagnosed with PTSD, while 19 were PTSD-. All analyses are Spearman’s Rank-Order correlations (*ρ < 0.05, **ρ < 0.01).

We then determined which individual symptoms correspond with myelin levels in the hippocampus, amygdala, and corpus callosum. We found that amygdala and hippocampal myelin estimates displayed divergent relationships with respect to individual PTSD symptoms. Specifically, hippocampal myelin estimates positively correlated with place avoidance (“Avoidance of activities, places, people”), mood (“Diminished interest in activities,” “Restricted affect”), sleep disruption, irritability, and concentration. In contrast, amygdala myelin positively correlated with emotional avoidance (“Avoidance of thoughts, feelings, conversations”), mood (“Diminished interest in activities”), psychological and physiological distress at exposure to cues, and startle. Interestingly, corpus callosum myelin estimates showed no significant relationships to any CAPS measures, a finding consistent with our work in rats.

In sum, hippocampal myelin in trauma-exposed men positively correlated with individual outcomes relating to place avoidance and hyperarousal, while amygdala myelin correlated with individual outcomes surrounding cue-associated distress, emotion, and startle. Both regions correlated with aspects of mood, and the corpus callosum, the brain’s largest white matter tract, showed no relationship to any aspects of PTSD symptomology. Together, these findings mirror several aspects of our work in rats. They provide clinical relevance to our findings and point to future avenues of investigation.

## Discussion

This study was motivated by the unmet critical need to understand the neural mechanisms that drive behavioral variation in response to acute, traumatic stress. Here, using a translational design, we identified region-specific gray matter oligodendrocytes and myelin as correlative indicators and potential contributors to anxiety-like behavior and fear learning after stress. We used a multimodal method of quantifying behavior to demonstrate that acute, severe stress yields individual variation in anxiety-like and fear behavior in rats. We found that persistent increases in place avoidance were positively correlated with dentate gyrus oligodendrocyte and myelin measures, while individual outcomes of contextual fear learning were positively correlated with myelin measures in the amygdala and spatial-processing subregions of the hippocampus. Together, these findings suggest that oligodendrocytes and myelin in these regions correspond with differential susceptibility to stress. Indeed, driving oligodendrogenesis in the dentate gyrus was sufficient to enhance avoidance and startle behavior in the absence of a stressor. Similarly, MRI-based estimates of gray matter myelin in trauma-exposed veterans positively correlated with PTSD symptomology in a region-specific manner: hippocampal myelin corresponded with place avoidance, and amygdala myelin corresponded with cue-associated fear. These findings provide a novel biological framework to understand how individual variance in persistent sensitivity to traumatic stress arises.

Traumatic stressors can be acute events that trigger long-lasting changes to the brain and behavior; however, the biological factors contributing to whether an individual develops mood and anxiety disorders after trauma remain poorly characterized (69). Modeling exposure to acute, severe stress is critical to understanding these mechanisms of susceptibility. In this study, we utilized a translational rat model in which animals were exposed to 3 hours of immobilization with simultaneous exposure to fox odor; this is an inescapable stressor that had been shown to be more severe than immobilization alone (57). Here, serum corticosterone was elevated throughout the duration of the stressor, and animals displayed significant weight loss by the day after stress, suggesting that this paradigm elicits a robust stress response.

Stress-induced changes to behavior may be isolated to a single domain or coordinated across multiple domains (5,64,70). To evaluate rat behavior after acute stress in a way that models complex and hypervigilance behavior, fear learning, and fear extinction. We found that the behavioral domain of avoidance clustered with exploration after acute stress, but not at baseline. This shift in behavioral signatures may speak to the nature of the open field test, which is often used as a means of quantifying either general exploratory locomotion or anxiety-like behavior (avoidance of the center). Our results suggest that under baseline conditions, open field behavior does not reflect anxiety-like behavior as measured by the elevated plus maze or light-dark box. However, after stress, avoidance of the center of the open field correlates well with avoidance measurements, hence serving as an assay of approach/avoidance conflict. Interestingly, we did not find strong correlations between avoidance, fear conditioning, or startle measures in stress-exposed animals, suggesting that, at least at the group level, this stress paradigm induces orthogonal behavioral effects with separable underlying neural mechanisms.

In this work, we developed composite cutoff behavior scoring to characterize behavioral profiles in rats. This method, and the Cutoff Behavioral Criteria from which it was adapted, does not rely upon absolute values of behavior, but instead applies binary scoring after application of a cutoff; summing scores from many measures and assays then allows for the identification of animals that display sensitive, yet consistent, anxiety-like behavior. This method is modeled upon human clinical standards and previous work in rodents (48,71,72). Similar thresholding techniques have been used to define distinct behavioral phenotypes in animals exposed to acute or chronic stress (64, 73). In our study, this method revealed that one week after acute stress exposure, animals display a spectrum of scores, from minimally affected to highly affected. Our approach models the continuous nature of stress effects and provides another means with which to investigate the physiological, neural, and glial factors that correspond and contribute to the continuum of anxiety-like behavior.

A growing number of studies implicate myelin and oligodendrocytes in behavioral outcomes associated with stress (10,13,47). Here, we showed that two weeks after an acute stressor, oligodendrocytes and myelin in the dentate gyrus of the hippocampus corresponded with outcomes of avoidance behavior on both a group and individual level. This relationship was specific to the dentate gyrus and was not present in the amygdala, corpus callosum, or CA regions of the hippocampus. In contrast, greater myelin content in the amygdala and CA2/3 regions of the hippocampus corresponded with greater contextual fear memory. These associations are fitting, as the hippocampus has been implicated in approach/avoidance conflict and anxiety (74–76), while the amygdala and CA hippocampal subregions contribute to associative fear learning and spatial memory, respectively (77, 78). A question that then arises is whether the correlation of oligodendrocytes to behavioral outcomes is merely an epiphenomenon (implying the potential to serve as a biomarker for vulnerability to stress-induced anxiety) or whether increased oligodendrogenesis plays a driving role in the pathogenesis of stress-induced anxiety outcomes (implying the potential to also serve as a therapeutic target). Here, we first showed that dentate gyrus oligodendrocytes born in the timeframe surrounding stress correspond to anxiety-like behavior outcomes at the group level. This may suggest that oligodendrogenesis plays a role in determining stress outcomes, whether via heightened baseline oligodendrogenesis, peri-traumatic oligodendrogenesis, or enhanced oligodendrocyte survival. To begin to parse these hypotheses, we conducted a gain-of-function experiment in which we ectopically overexpressed the oligodendrocyte transcription factor Olig1 in the dentate gyrus of the hippocampus. We demonstrated that heightened oligodendrocytic drive in the dentate gyrus is sufficient to bring about increased avoidance and startle behavior. This moves our findings beyond correlational work and suggests that hippocampal oligodendrogenesis can contribute to anxiety-like behavior. Of note, we chose a constitutive promoter that could target multiple cell types, but a clear limitation of this approach is the non-specificity of transgene expression. In addition, loss-of-function experiments will be needed to address the question of the necessity of oligodendrogenesis for stress-induced anxiety-like behavior, which may ultimately contribute to the development of pharmacological interventions for trauma-induced disorders. Lastly, we focused our gain-of-function experiment on the hippocampus, but future work should seek to conduct additional gain- and loss-of-function experiments in sites such as the amygdala. Nonetheless, this experiment further adds to the growing body of work demonstrating that adaptive myelination is not a passive epiphenomenon but a contributing factor to experience-dependent plasticity and neural circuit function.

This study adds an important dimension to a growing body of literature demonstrating that oligodendrocytes outside of white matter tracts are associated with behavior and stress. Notably, several studies show that chronic stress paradigms decrease myelin in regions such as the PFC or in white matter tracts (24, 25). However, *increases* in oligodendrogenesis and myelination can be induced by experience and neural activity in regions such as premotor cortex, PFC, hippocampus, retrosplenial cortex, and others (14,20,26,32). Moreover, inhibiting oligodendrogenesis and the formation of new myelin can impair motor learning, fear recall, and social behavior, suggesting that increases in myelination can alter behavior (14,21,26,45). Our work expands upon such results by detailing additional gray matter regions in which oligodendrocytes may influence behavior; specifically, we implicate oligodendrocytes and increased myelin in the hippocampus and amygdala in anxiety-like phenotypes following stress exposure. In addition, our data support and expand upon a recent study that utilized microarray gene expression data from animals exposed to acute stress, which demonstrated that hippocampal oligodendrocyte proportions are elevated in animals displaying PTSD-like behavioral responses (79). Here, we find similar increases in oligodendrocyte density in the hippocampus of animals with high composite anxiety-like behavior scores. We further demonstrate a correlative relationship between behavioral outcomes and hippocampal oligodendrocyte content, we provide evidence suggesting a causal relationship between the two, and we demonstrate the clinical relevance of this phenomenon. Ultimately, our findings suggest a role for gray matter oligodendrocytes and myelin in selective susceptibility to stress-induced anxiety, adding to the growing hypothesis that myelin in the central nervous system contributes to the pathophysiology of psychiatric conditions.

Importantly, we employed a translational approach in this study; our results from rats informed our experiments with humans and vice versa. We previously showed that hippocampal myelin positively correlates with overall PTSD symptom severity (46). Here, we have now modeled this relationship in a controlled rat experiment and extended it to show that oligodendrocytes and myelin in the hippocampus as well as the amygdala correspond in a region-specific manner to individual outcomes of avoidance, startle, and fear-like behavior. Given these findings in rats, we then tested whether region-specific myelin estimates corresponded to individual components of PTSD symptom profiles in a group of trauma-exposed veterans. We found that estimates of hippocampal myelin correlated with place avoidance, sleep disturbances, and concentration. In contrast, amygdala myelin correlated with cue-associated distress, emotional avoidance, and exaggerated startle. This adds a comparative and translational perspective to our findings in rats. For example, the relationships between hippocampal myelin and avoidance and between amygdala myelin and contextual fear were observed in both our rat and our human data. These parallels imply that such relationships between gray matter myelin and behavior are evolutionarily conserved and species-independent. In addition, they emphasize that relationships between myelin and individual domains of symptoms and behavior are region-specific. Ultimately, this study represents an example of an expressly translational investigation in which similar brain regional biomarkers are correlated with nominally analogous behavioral indices across rats and humans, and the findings illustrate important species similarities and differences that can guide future translational work towards aspects of myelin-behavior relationships that are likely to be conserved.

The relationships observed in this study among oligodendrocyte numbers, myelination, and behavioral outcomes may reflect different biological scenarios. First, baseline variation in the density of oligodendrocytes and myelin may confer variable sensitivity to stress exposure. Alternatively, variability in physiological stress responses may differentially regulate oligodendrogenesis. Support for either hypothesis will require quantification of oligodendrocytes and myelin *in vivo* in a longitudinal design, utilizing methods such as MRI or temporally-controlled genetic techniques to label new myelin in a way that still allows for behavioral profiling. In addition, future work should seek to incorporate other hippocampal- and amygdala-specific behavioral tasks that show deficits in PTSD-related behavior, such as spatial navigation or cued fear learning (80, 81), in order to elucidate how region-specific myelin measures correspond to the full array of trauma-induced behavioral outcomes. Lastly, expanding this work to include additional regions of interest will be critical to developing a comprehensive, brain-wide model of how myelin relates to individual stress outcomes in both rats and humans.

Added to these unresolved considerations is the question of sex differences. Our human cohort, both here and in our previous publication, consisted solely of male veterans due to the relative scarcity of female veterans who met all study criteria. We therefore focused our rat work on males, and our results should not be generalized to females. Given the hypothesized sex differences of psychiatric and behavioral outcomes in women and female rodents after stress (82–84), future studies should seek to explore both the behavioral outcomes of female rats after acute, severe stress, as well as the relationships between stress-induced behavior and myelin in female rat and human populations.

This study ultimately presents important insight for the understanding of psychiatric disorders. Abnormalities in myelin and oligodendrocytes have been implicated in numerous disorders, including PTSD, depression, schizophrenia, autism, and Alzheimer’s disease (10,44,46,85–88). Our findings here contribute to the growing hypothesis that myelin in the central nervous system contributes to the pathophysiology of psychiatric conditions by suggesting a role for gray matter oligodendrocytes and myelin in selective susceptibility to stress-induced anxiety. While molecular phenotyping of tumor markers transformed the field of cancer diagnosis and treatment, the field of psychiatry is lagging far behind any promise of personalized medicine. By identifying regional relationships between myelin and behavior, our findings may ultimately lead to much needed imaging-based tools for mechanism-based patient stratification that will aid in identifying and treating the most vulnerable individuals.

## Supporting information

Correlation and T Test Statistics

## Acknowledgments

We thank Ellen Brooker, Linnea Delucchi, Brian Gallagher, Natalie Musick, David Seo, Chau Pham, Lydia Ross, Katerina Alexander, Anjana Krishnamurthy, Amalia Kaufer Shapira, Anya Fineman, Iris Hart, and Cory Berg for assistance with animal work and data collection. We also thank Kristen Berendzen for thoughtful edits to the manuscript. Slide imaging experiments were conducted at the Cancer Research Laboratory Molecular Imaging Center, supported by the UC Berkeley Biological Faculty Research Fund. We would like to thank Holly Aaron and Feather Ives for their microscopy training and assistance. We thank Mary West, Pingping He, and the UC Berkeley QB3 Cell & Tissue Analysis Facility / High-Throughput Screening Facility for lentiviral packaging.

## Author contributions

D.K. and K.L. devised experiments. K.L. performed experiments and analyses. L.L.C. conducted and analyzed human MRI. Y.K., A.A., K.H., L.P., D.M., V.R., R.M., C.T., and J.B. performed animal work, immunohistochemistry, and analysis. W.C. and S.M. provided oversight and support for statistical analyses. B.R.H. and S.H.W. supervised human PTSD analyses. T.C.N. and D.K. directed the project. K.L., Y.K., D.M., L.P., and D.K. wrote the manuscript.

## Data and materials availability

All data associated with this study are available in the main text or the supplementary materials.

## Supplementary Materials

Fig. S1: Measures from behavior profiling tests.

Fig. S2: Behavior cluster composition.

Fig. S3: Composite cutoff scoring.

Fig. S4: Relationships between brain measures and composite behavior scores in unexposed, control animals.

Fig. S5: Behavioral measures from avoidance tests in Lenti-GFP vs. Lenti-Olig1 animals.

Fig. S6: Longitudinal relationships between avoidance, startle, and fear conditioning behavior.

