## Supplemental Figures for "Region-specific, maladaptive, gray matter myelination is associated with differential susceptibility to stress-induced behavior in rats and humans"

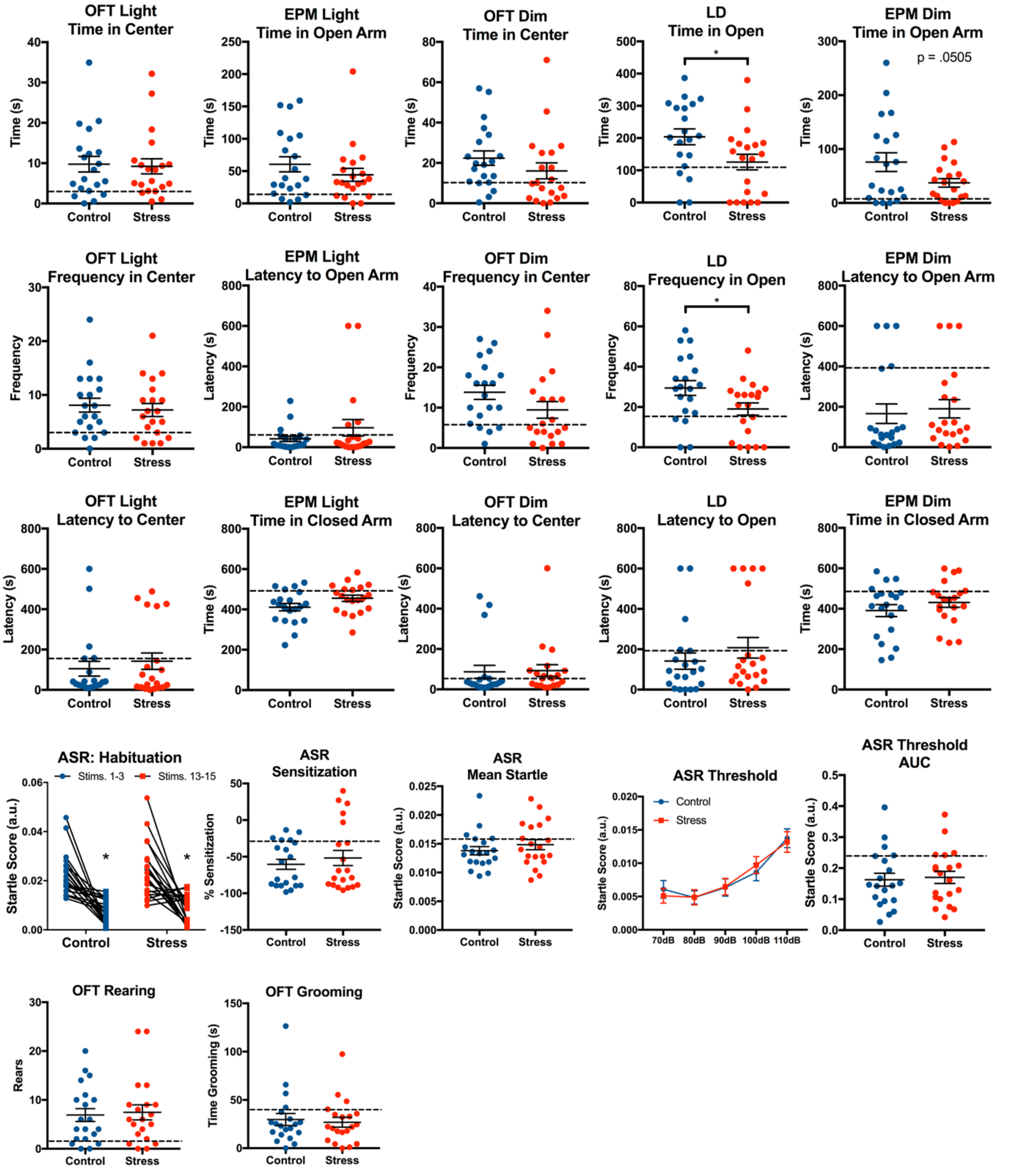


**Fig. S1: Measures from behavior profiling tests.** Raw values from each behavioral measure are shown. Dotted lines demarcate the 20th percentile (for measures of approach towards anxiogenic zones) or 80th percentile (for measures of avoidance of anxiogenic zones) of the control distribution that were used as cutoff criteria. a.u., arbitrary units. *$p < 0.05$

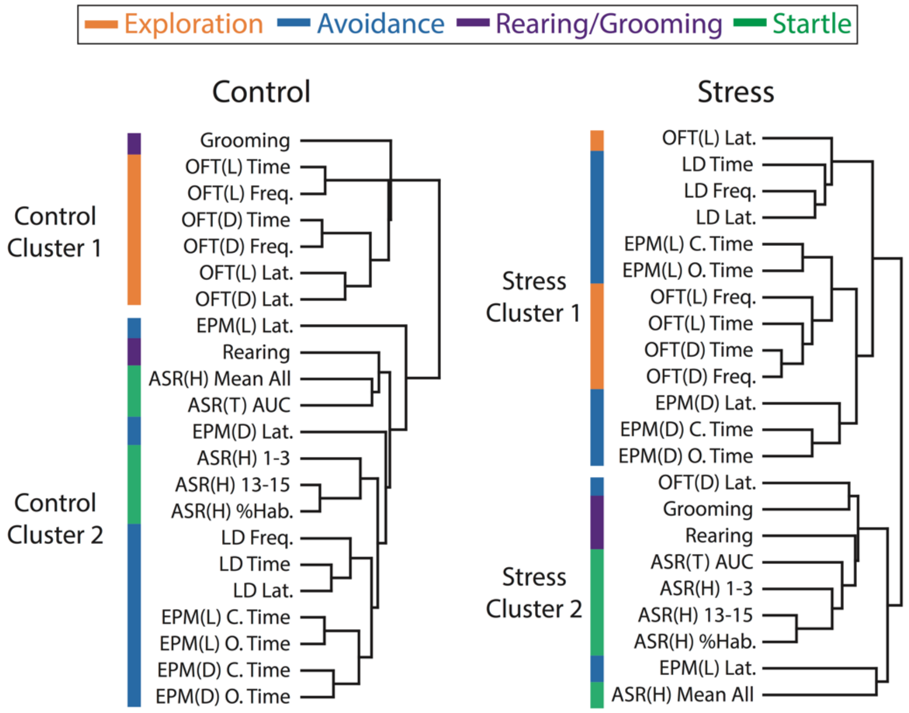


**Fig. S2: Behavior cluster composition.** Results of hierarchical clustering of all behavioral measures. OFT measures: Time, time in center; Freq., frequency in center; Lat., latency to center. EPM measures: Lat., latency to open arm; C. Time, time in closed arm; O. Time, time in open arm. LD measures: Time, time in open zone; Freq., frequency in open zone; Lat., latency to open zone. ASR(H), ASR habituation phase: Mean All, mean startle to all 15 110 dB stimuli; 1-3, average startle to stimuli 1-3; 13-15, average startle to stimuli 13-15; %Hab., percent habituation. ASR(T) AUC, ASR threshold phase area under the curve.

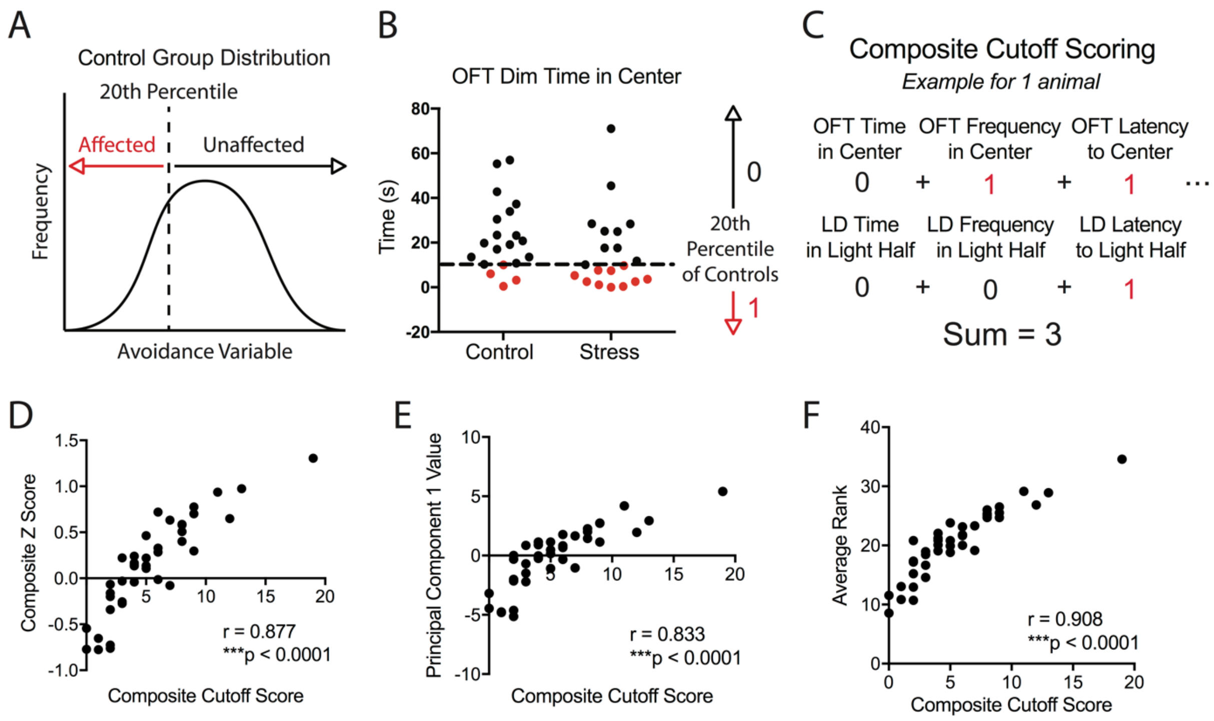


**Fig. S3: Composite cutoff scoring. (A)** Defining the cutoff criterion: For each behavioral measure, the cutoff is defined as the 20th percentile of the control group distribution. Values falling below the cutoff criterion are assigned an "Affected" score. **(B)** Example of defining affected values. Values falling below the 20th percentile of the control distribution are scored as affected. **(C)** All affected scores are summed across the avoidance tests. **(D)** Pearson correlation of composite cutoff scores to composite Z scores (Pearson correlation: r = 0.88, $p < 0.0001$; n = 40). **(E)** Pearson correlation of composite cutoff scores to principal component 1 values of a principal component analysis (Pearson correlation: r = 0.83, $p < 0.0001$; n = 40). **(F)** Pearson correlation of composite cutoff scores to the average of an animal's rank within each test (Pearson correlation: r = 0.91, $p < 0.0001$; n = 40).

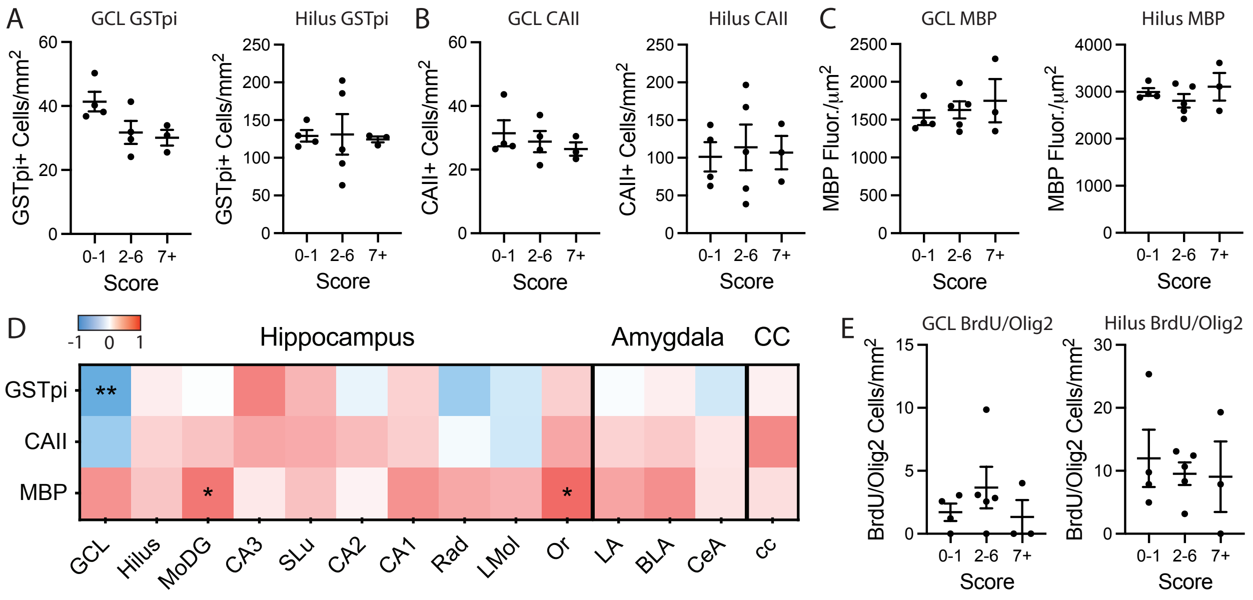


**Fig. S4: Relationships between brain measures and composite behavior scores in unexposed, control animals. (A)** Dentate gyrus GSTπ measures from animals with low (0-1), mid-range (2-6), and high (7 or more) composite behavior scores. All data are presented as individual values with mean ± SEM. **(B)** Dentate gyrus CAII measures. **(C)** Dentate gyrus MBP measures. **(D)** Correlation matrix of composite behavior scores to oligodendrocyte and myelin measures from a subset of animals (Pearson correlations: n = 11-12; one high outlier removed from each of GCL GSTπ and CeAmy GSTπ datasets; one high outlier removed from GCL CAII dataset; 3 low outliers removed from MoDG CAII dataset). **(E)** Dentate gyrus BrdU/Olig2 colabeling measures.

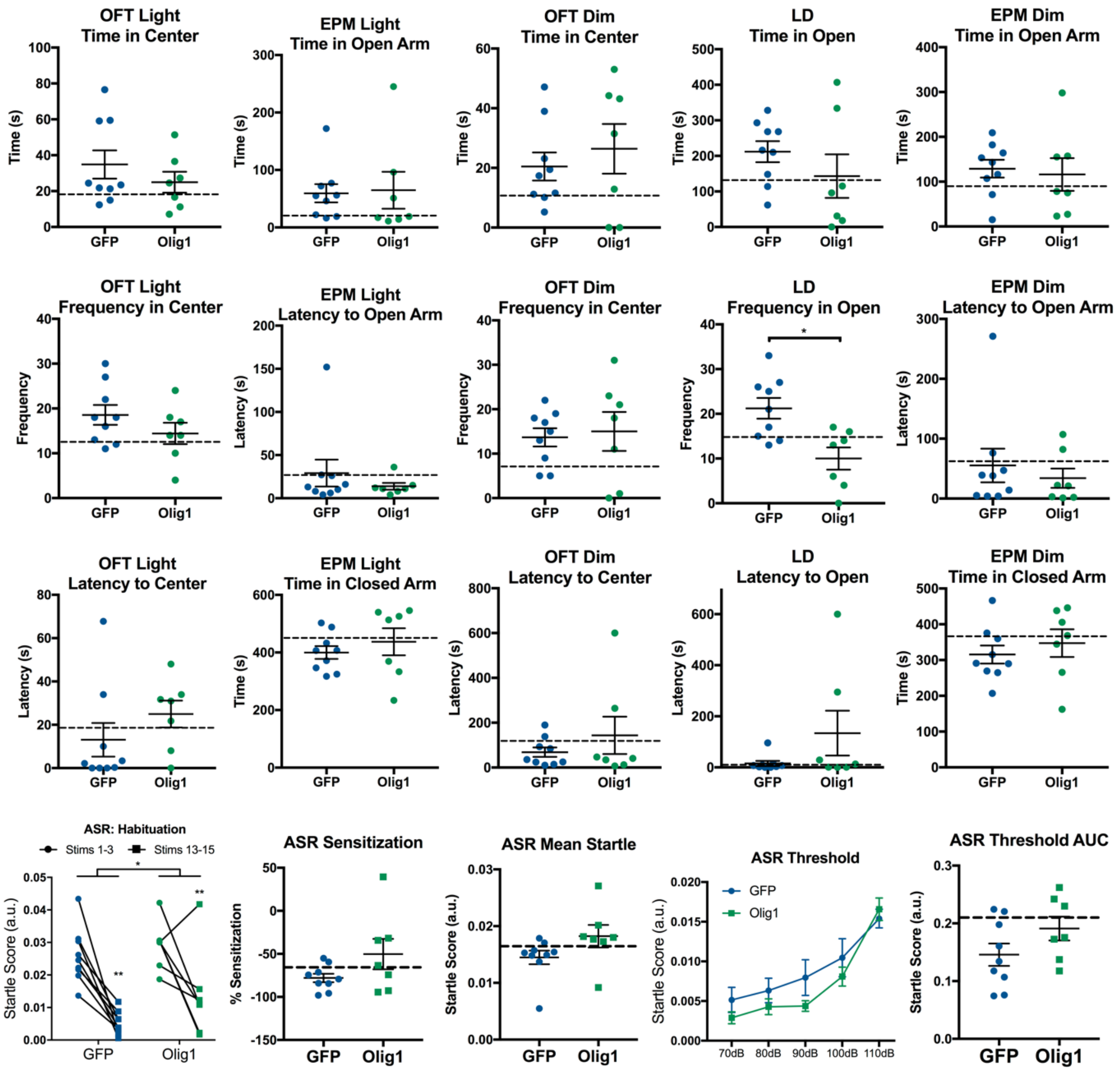


**Fig. S5: Behavioral measures from avoidance tests in Lenti-GFP vs. Lenti-Olig1 animals.** Raw values from each behavioral measure are shown. Dotted lines demarcate the 20th percentile (for measures of approach towards anxiogenic zones) or 80th percentile (for measures of avoidance of anxiogenic zones or high startle activity) of the control distribution that were used as cutoff criteria for the composite behavioral scores.

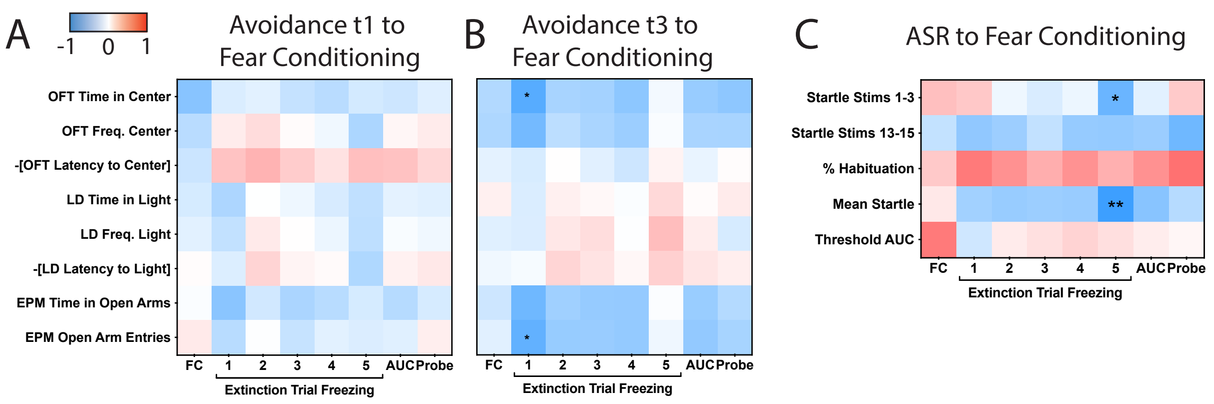


**Fig. S6: Longitudinal relationships between avoidance, startle, and fear conditioning behavior. (A)** Correlation matrix of avoidance measures at time point 1 (t1) to fear conditioning measures. FC, fear conditioning; AUC, area under the curve; Pearson correlations: n = 12). **(B)** Correlation matrix of avoidance measures at t3 to fear conditioning measures (Pearson correlations: n = 12). **(C)** Correlation matrix of acoustic startle response (ASR) measures to fear conditioning measures (Pearson correlations: n = 12). **p* < 0.05.
